# DeepGenePrior: A deep learning model to prioritize genes affected by copy number variants

**DOI:** 10.1101/2022.08.22.504862

**Authors:** Zahra Rahaie, Hamid R. Rabiee, Hamid Alinejad-Rokny

## Abstract

The genetic etiology of neurodevelopmental disorders is highly heterogeneous. They are characterized by abnormalities in the development of the central nervous system, which lead to diminished physical or intellectual capabilities. Determining which gene is the driver of disease (not just a passenger), termed ‘gene prioritization,’ is not entirely known. In terms of disease-gene associations, genome-wide explorations are still underdeveloped due to the reliance on previous discoveries when spotting new genes and other evidence sources with false positive or false negative relations. This paper introduces DeepGenePrior, a model based on deep neural networks that prioritizes candidate genes in Copy Number Variant (CNV) mediated diseases. Based on the well-studied Variational AutoEncoder (VAE), we developed a score to measure the impact of the genes on the target diseases.

Unlike other methods that use prior data on gene-disease associations to prioritize candidate genes (using the guilt by association principle), the current study exclusively relies on copy number variants. Therefore, the procedure can identify disease-associated genes regardless of prior knowledge or auxiliary data sources. We identified genes that distinguish cases from disorders (autism, schizophrenia, and developmental delay). A 12% increase in fold enrichment was observed in brain-expressed genes compared to previous studies, while 15% more fold enrichment was found in genes associated with mouse nervous system phenotypes. We also explored sex dimorphism for the disorders and discovered genes that overexpress more in one gender than the other. Additionally, we investigated the gene ontology of the putative genes with WebGestalt and the associations between the causative genes and the other phenotypes in the DECIPHER dataset. Furthermore, some genes were jointly present in the top genes associated with the three disorders in this study (i.e., autism spectrum disorder, schizophrenia, and developmental delay); namely, deletions in *ZDHHC8*, *DGCR5*, and *CATG00000022283* were common between them. These findings suggest the common etiology of these clinically distinct conditions.

With DeepGenePrior, we address the obstacles in existing gene prioritization studies. This study identified promising candidate genes without prior knowledge of diseases or phenotypes using deep learning.

## Introduction

Neurodevelopmental disorders (NDDs) [1] are a group of disorders that affect the development of the nervous system. A dysfunctional brain can influence memory, emotion, and learning ability. Deleterious variations such as deletions in 16p11.2 [2–4] and duplications in 15q3 [5, 6] are well-studied loci associated with autism (a type of NDD). Genetic factors related to autism include *TBX1* (in the 22q11.2 deletion syndrome – a transcription factor involved in the regulation of development), *SHANK3* (a synaptic scaffolding gene), *NLGN4* (a neuroligin gene), *PCDH10* (a protocadherin gene, a subfamily of the cadherin superfamily) and *NHE9* [7, 8]. Others include *NRXN1, SHANK2, CNTN4,* and *CNTNAP2*. In addition, *DPYD* and *DPP6* are candidate genes for autism, as are *RFWD2*, *NLGN1*, *ASTN2*, *SYNGAP1*, and *DLGAP2*, as well as *DDX53-PTCHD1* [9].

NDD includes schizophrenia (SCZ) as another disorder; CNVs disrupt several genes in SCZ, including *TBX1* (common in autism), *ERBB4* (type I receptor tyrosine kinase subfamily, encodes a receptor for NDF/heregulin), *SLC1A3* (solute carrier family 1, glutamate transporter, member 3), *RAPGEF4* (rap nucleotide exchange factor) and *CIT* (a neuronal Rho-target gene) [7, 8]. 7q11.2 and 15q13.3 were reported by [10] to be associated with SCZ. In the SCZ, a large (3 Mb) deletion on chromosome 22q11.21 is a significant risk factor [9]. Besides, other loci, including deletions at 1q21.1, deletions at 3q29, duplications of 16p11.2, deletions at 15q13.3, exonic deletions at 2p16.3, and duplications at 7q36.3, were also reported [9].

For the developmental delay, deletions in 1q24 (*FMO* group of genes and *DNM3*), 2q33.1 (*SATB2*), and 2p16.1 (*NRXN1*) are some of the well-known variations. [11]

Research on the genetics of diseases has significant implications for diagnosing, treating, and developing drugs for them. The genetic etiology of neurodevelopmental disorders can provide substantial insight into effective prevention and treatment methods. Identifying the genes that most likely contribute to a disease or phenotype is a process known as gene prioritization.

The prioritization of genes has been based on several pieces of evidence. Gene-disease associations are classified into five categories [12]: functional, cross-species, same-compartment, mutation, and textual. Molecule interactions are examined in the first category (such as [13]). In the second category, homolog genes causing similar phenotypes are explored in other organisms (such as [14]). Same-compartment evidence focuses on the fact that the gene participates in known disease-associated pathways or compartments (for instance, the cell membrane or nucleus) (like [15]). Mutation evidence relies on SNPs and structural variations (which is also the focus of this study) (such as [16]). PubMed and other online collections can be used to collect text evidence (such as [17]).

Gene prioritization methods are reviewed in [18–21]; the methods can be categorized into two groups, statistical and machine learning. To determine whether a gene is associated or not, the first group mainly uses hypothesis testing, such as exact tests like Fisher’s or permutation tests. In several studies (such as [22]), p-value fallacies have been discussed, such as distributional assumptions, not assigning probabilities to hypotheses, limitations in data collection, and misleading results. In [3], power loss and dependent values are discussed in detail as other criticisms of marginal p-values.

Afterward, machine learning methods require some seed data (in this case, genes that implicitly characterize the disorder) [18] and a similarity metric (to assess how far candidate genes differ from known ones); They then determine which candidate gene is similar or has an association with the seed. There is a fundamental principle behind this category, known as ‘guilt by association,’ which refers to genes discovered that are related to already known genes. Some issues arise with this approach. In other words, we cannot expect to find a novel gene association that does not have any relation to the previous ones; In addition, we cannot discover the genes of a genetic disease that we do not know anything about [23, 24].

A flawless gene prioritization solution is hindered by the issues discussed above. In this study, we propose DeepGenePrior as a deep learning architecture for gene prioritization to avoid these issues. It combines a well-studied autoencoder architecture with a variational learning framework. Variational AutoEncoder [25, 26] (shortened as VAE, which is used in the text) is the stochastic variant of the autoencoder. Our first step is to measure the overlaps between the CNVs and the genes in the training set. Controls and cases have zero and one CNV labels, respectively. We then train the network of neurons with all CNVs of cases and controls for all three diseases. Next, we fine-tune it with the CNVs of the target disease. As a final step, we built a score for every gene using the network weights and prioritized them. Fig. 1 summarizes the method.

**Figure 1:**
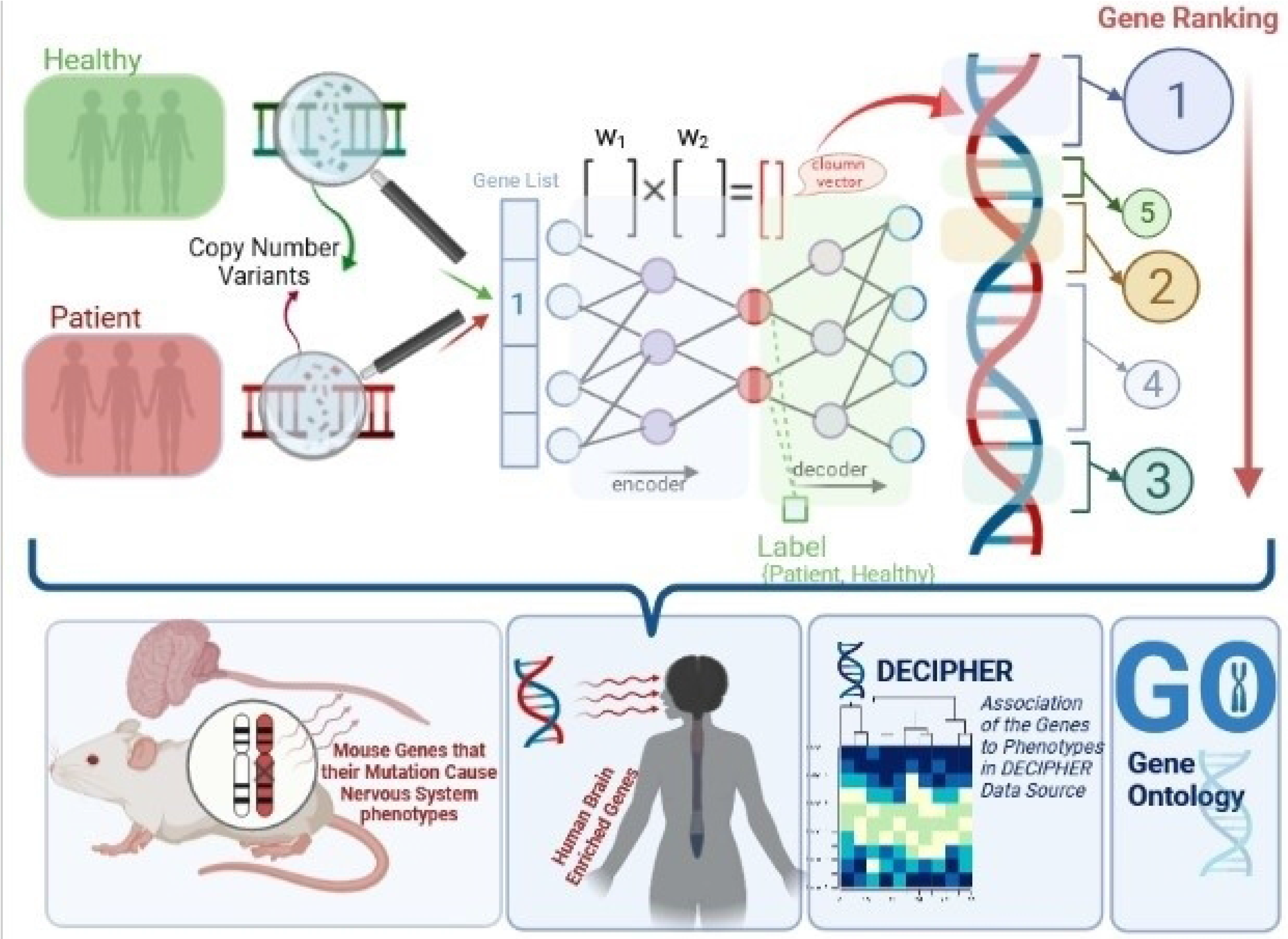
Graphical Abstract; the Summary of the Method and Analyses of the Results.

The method addresses the gaps in the previous studies. There are several advantages to this method, including that it is not based on unreal theoretical assumptions (such as hypothesis tests), it does not require seed data (to be used by guilt by association), and it does not rely on networks with false positive relations (like protein-protein networks). Additionally, pretraining uses all available data.

Using CNVs from neurodevelopmental disorders, we benchmark our method against major tools. We find significantly mutated genes; interestingly, our method finds genes that are 12% more enriched in brain expression than other tools. Furthermore, we compared the detected genes with those whose mutations cause nervous system phenotypes in mice. Compared to other methods, our results were 15% more enriched.

In addition, we examined genes that were exclusively overrepresented in one gender. Besides, we analyzed the relationships between the detected genes and the various phenotypes in the DECIPHER [27] data source, as well as the gene ontology of the putative genes. Three genes were common in the top genes associated with the three diseases; deletions in *ZDHHC8*, *DGCR5*, and *CATG00000022283* were common between the three disorders; according to the literature, defects found in *ZDHHC8* can be linked to the susceptibility to schizophrenia. Also, deletions in *CYFIP1*, *PRODH*, *XXBAC*, *B444P24*, *LINC00896*, *ZDHHC8*, *AC006547*, *NIPA2*, *RTN4R*, *NIPA1*, and *TUBGCP5* are found to be associated with schizophrenia and developmental delay.

The following section describes the algorithm, the data, the experiments, and the analysis of the results. Finally, the last section presents the conclusions and future works. Table 1 provides the list of abbreviations.

**Table 1:** List of short abbreviations used in the text.

| Abbreviation | Long Word |
| --- | --- |
| CNV | Copy Number Variant |
| VAE | Variational Autoencoder |
| DD | Developmental Delay |
| SCZ | Schizophrenia |
| ASD | Autism Spectrum Disorder |
| NDD | Neurodevelopmental Disorders |
| AUC | Area Under the Curve |
| DECIPHER | Database of Chromosomal Imbalance and the phenotype in Humans Using ENSEMBL Resources |
| WebGestalt | WEB-based Gene Set Analysis Toolkit |

## Results

### Prioritization of the Genes in Neurodevelopmental Disorders

A deep learning model was used to identify the genes underlying a disorder. This model aims to uncover the genes contributing to the target disease and the underlying relationship patterns. Based on the copy number variants of all cases and controls, we train a network, then use the model weights to calculate scores.

Our CNV datasets come from three sources: autism spectrum disorder [28], developmental delay [11], and schizophrenia [29]. A UCSC Lift Genome Annotations tool was used to convert all CNVs to hg19, and NCBI remap tools were used to confirm the locations of all CNVs. Those CNVs smaller than 1KBp were eliminated. Tables 8, 10, and 11 show the top 40 genes for each disorder with their corresponding p-values (with Fisher’s exact test shown in Table 9).

**Table 2:**
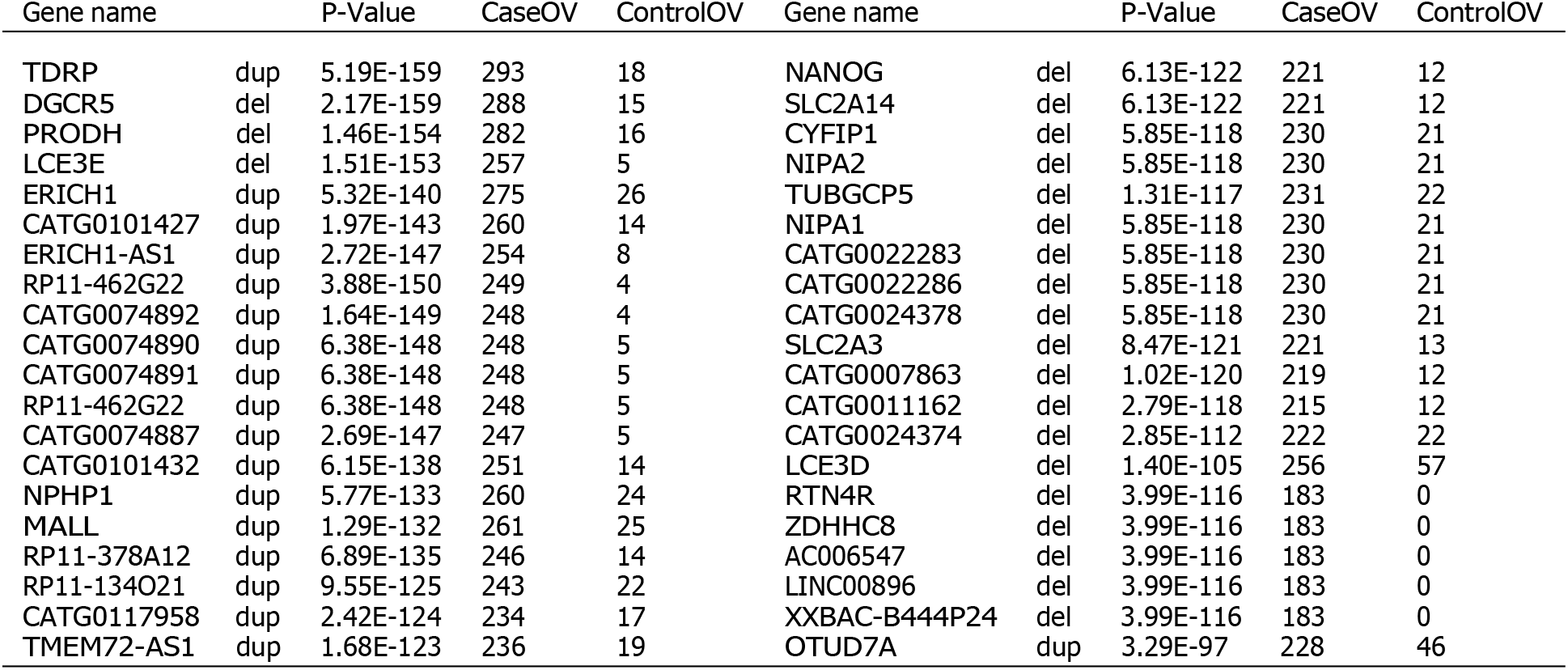
Top 40 genes related to developmental delay.

**Table 3:** Contingency table for Fisher’s exact test.

|  |  |
| --- | --- |
| # of case samples<br>overlapped with the gene | # of case samples not<br>overlapped with the gene |
| # of control samples<br>overlapped with the gene | # of control samples not<br>overlapped with the gene |

**Table 4:**
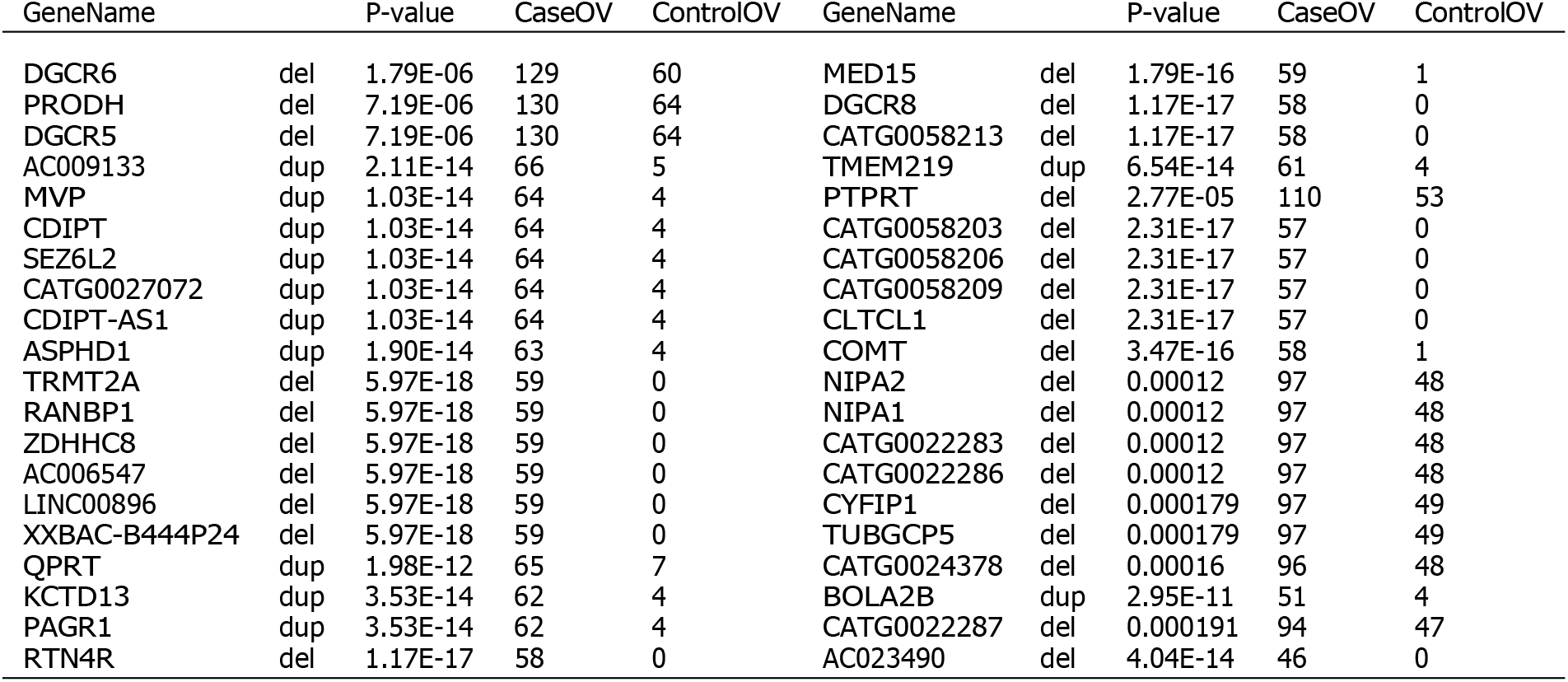
Top 40 genes related to schizophrenia.

**Table 5:** Top 40 genes related to ASD.

| Gene Name |  | P Val | CaseOV | ControlOV | Gene Name |  | P Val | CaseOV | ControlOV |
| --- | --- | --- | --- | --- | --- | --- | --- | --- | --- |
| DGCR2 | del | 2.38E-44 | 420 | 3 | GABRA5 | dup | 9.82E-28 | 235 | 0 |
| ARVCF | del | 3.77E-44 | 418 | 3 | OCA2 | dup | 8.85E-28 | 234 | 0 |
| GNB1L | del | 3.77E-44 | 418 | 3 | CATG00000022283 | del | 5.01E-08 | 263 | 32 |
| CATG00000058206 | del | 6.34E-44 | 417 | 3 | CATG00000022351 | dup | 2.34E-27 | 231 | 0 |
| COMT | del | 6.34E-44 | 417 | 3 | NRXN1 | del | 2.08E-12 | 249 | 19 |
| ZDHHC8 | del | 6.34E-44 | 417 | 3 | LINC00624 | del | 3.24E-15 | 241 | 13 |
| HIRA | del | 6.34E-44 | 417 | 3 | XXBAC-B135H6 | del | 1.66E-21 | 229 | 4 |
| TBX1 | del | 6.34E-44 | 417 | 3 | BCL9 | del | 1.01E-16 | 234 | 10 |
| CDIPT | del | 4.15E-28 | 387 | 15 | CHD1L | del | 1.01E-16 | 234 | 10 |
| SEZ6L2 | del | 4.15E-28 | 387 | 15 | FMO5 | del | 1.01E-16 | 234 | 10 |
| ASPHD1 | del | 4.15E-28 | 387 | 15 | ACP6 | del | 1.01E-16 | 234 | 10 |
| KCTD13 | del | 4.15E-28 | 387 | 15 | CATG00000092640 | del | 5.84E-17 | 141 | 0 |
| CATG00000027072 | del | 4.15E-28 | 387 | 15 | CATG00000058020 | del | 1.96E-15 | 142 | 1 |
| CDIPT-AS1 | del | 4.15E-28 | 387 | 15 | RFC2 | del | 1.94E-15 | 141 | 1 |
| ALDOA | del | 4.04E-28 | 386 | 15 | WBSCR22 | del | 9.53E-17 | 140 | 0 |
| FAM57B | del | 4.04E-28 | 386 | 15 | GTF2I | del | 9.53E-17 | 140 | 0 |
| CHRNA7 | del | 1.75E-26 | 239 | 1 | STX1A | del | 9.53E-17 | 140 | 0 |
| DGCR5 | del | 3.40E-21 | 244 | 6 | EIF4H | del | 1.94E-15 | 141 | 1 |
| GABRB3 | dup | 9.82E-28 | 235 | 0 | DNAJC30 | del | 9.53E-17 | 140 | 0 |
| OTUD7A | del | 2.64E-26 | 236 | 1 | VPS37D | del | 9.53E-17 | 140 | 0 |

**Table 6:** Brain-enrichment comparison, coding genes.

|  | Deletion | Duplication |
| --- | --- | --- |
| Genome-wide Expectation | 34.83% | 34.83% |
| Coe et al. [11] | 35.3% | 34.9% |
| Cooper et al. [33] | 35.1% | 33.7% |
| Our Result (DD) | 52.3% | 65.5% |
| Plink [34] | 35.1% | 36.3% |
| Our Result (SCZ) | 47.3% | 60% |
| SNATCNV (ASD) [28] | 40.4% | 45.5% |
| Our Result (ASD) | 48.4% | 61% |

**Table 7:** Brain-enrichment comparison, noncoding genes.

|  | Deletion | Duplication |
| --- | --- | --- |
| Genome-wide Expectation | 25.67% | 25.67% |
| Coe et al. [11] | 32.4% | 31.7% |
| Cooper et al. [33] | 33.4% | 31.7% |
| Our Result (DD) | 42.6% | 47.6% |
| Plink [34] | 28.3% | 33.9% |
| Our Result (SCZ) | 37.1% | 28.2% |
| SNATCNV (ASD)[28] | 33.6% | 45.2% |
| Our Result (ASD) | 37% | 46.5% |

Moreover, we investigate which genes are associated with three disorders and which are associated with two. Fig. 10 illustrates this. It suggests that these clinically distinct neurodevelopmental conditions are related in etiology.

**Figure 2:**
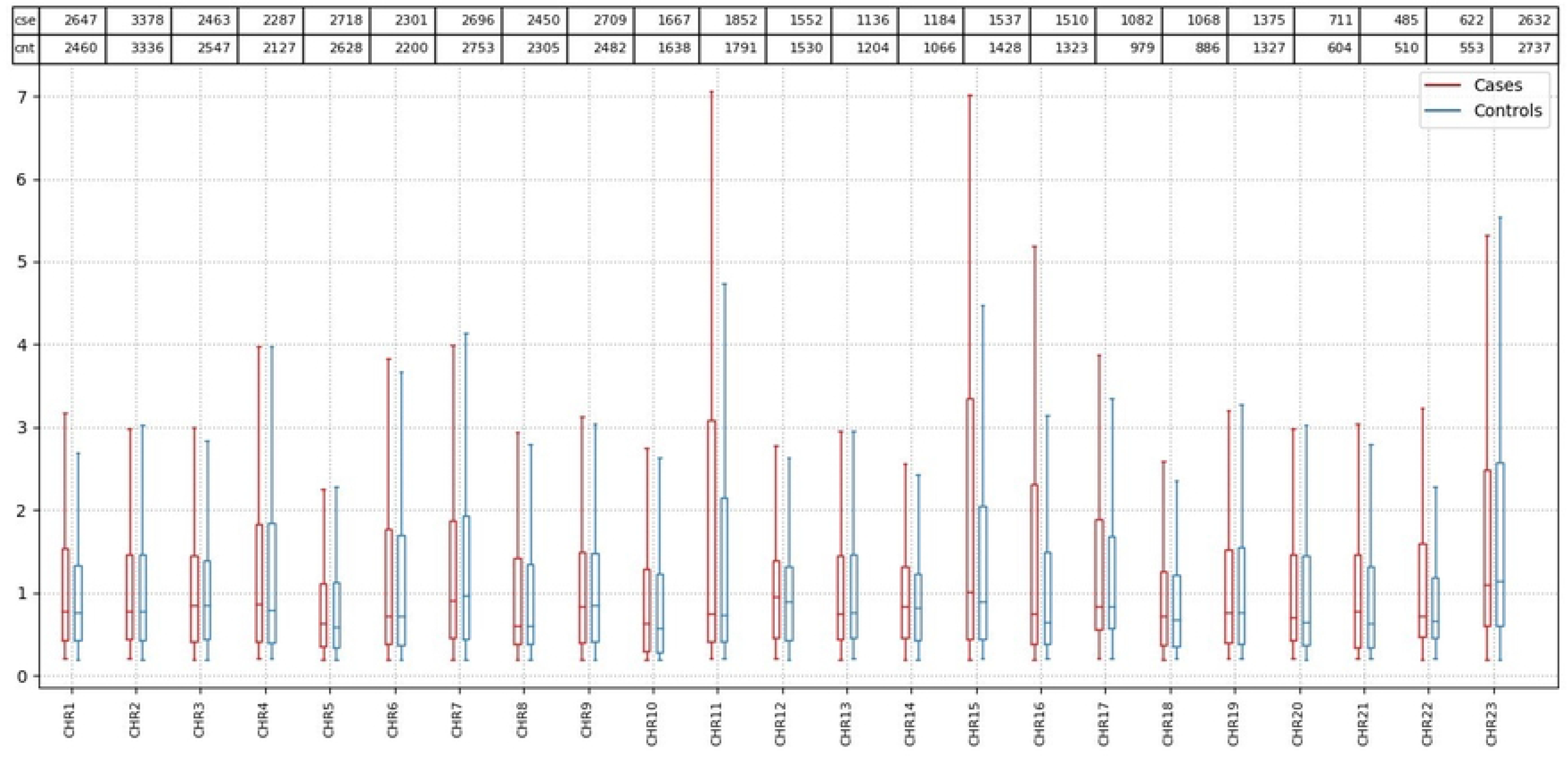
The common genes between two or three disorders ‘del’ is short for deletion.

**Figure 3:**
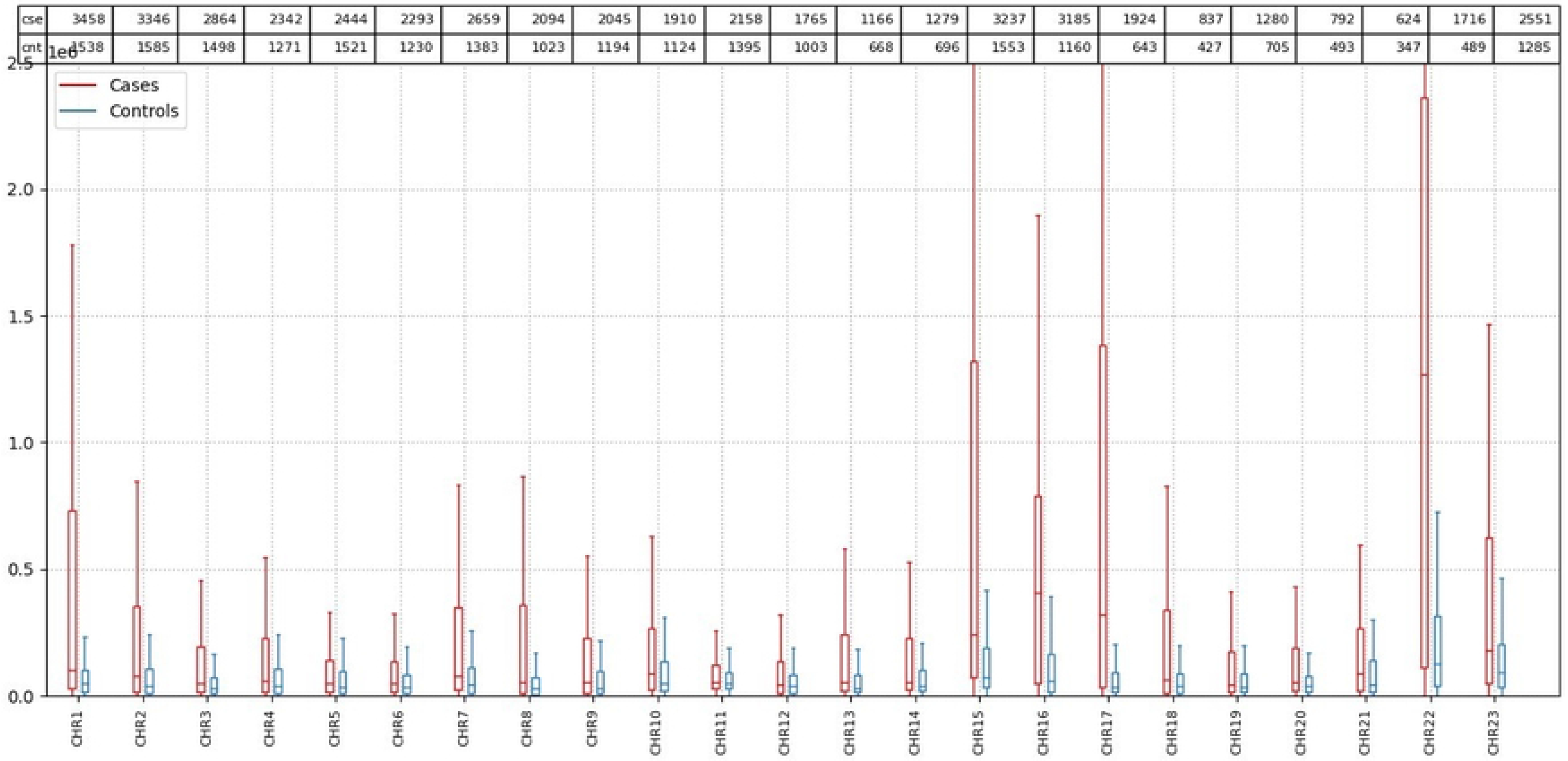
Decipher Phenotypes Frequency.

**Figure 4:**
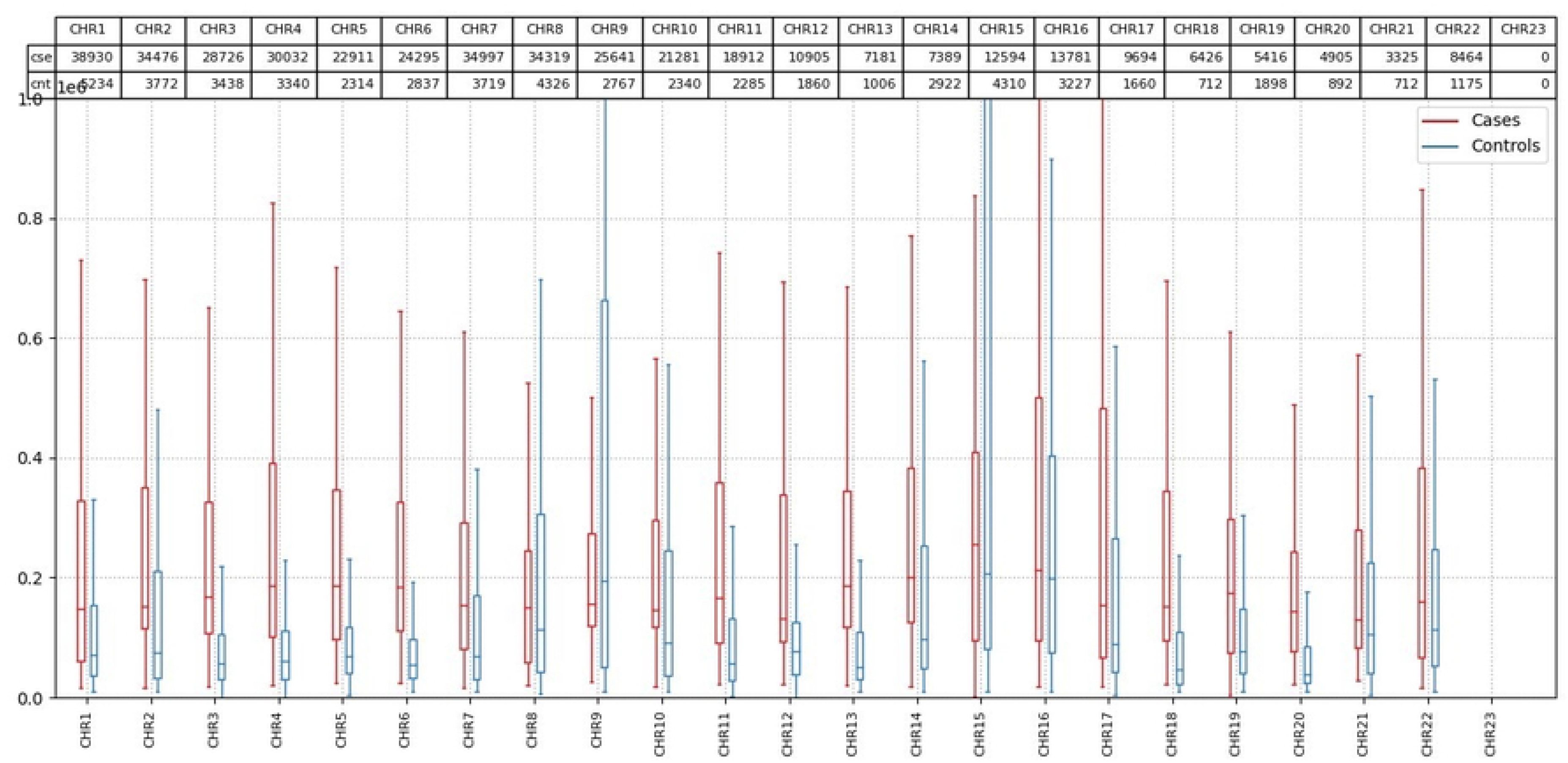
Heatmap for developmental delay.

**Figure 5:**
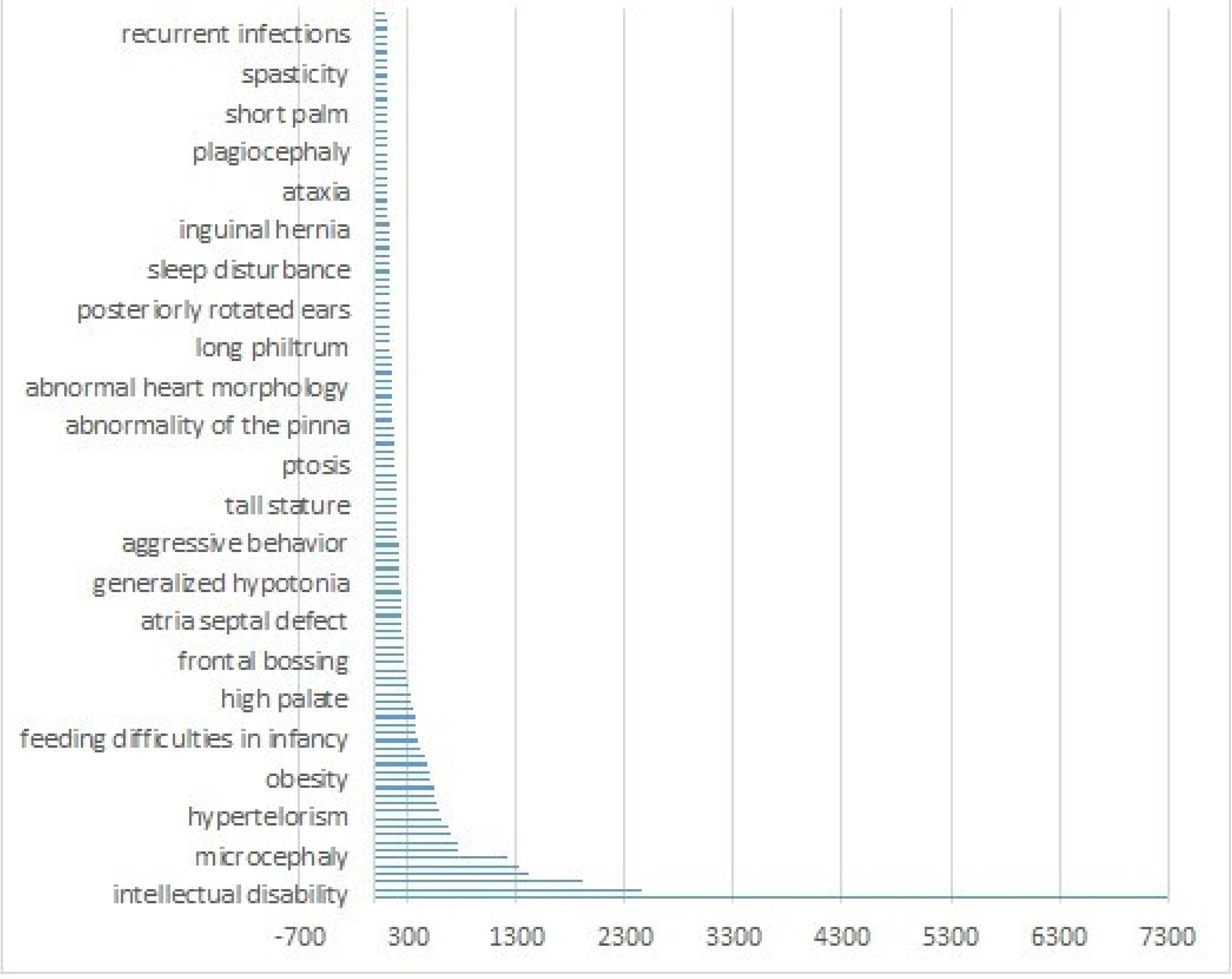
Heatmap for schizophrenia.

**Figure 6:**
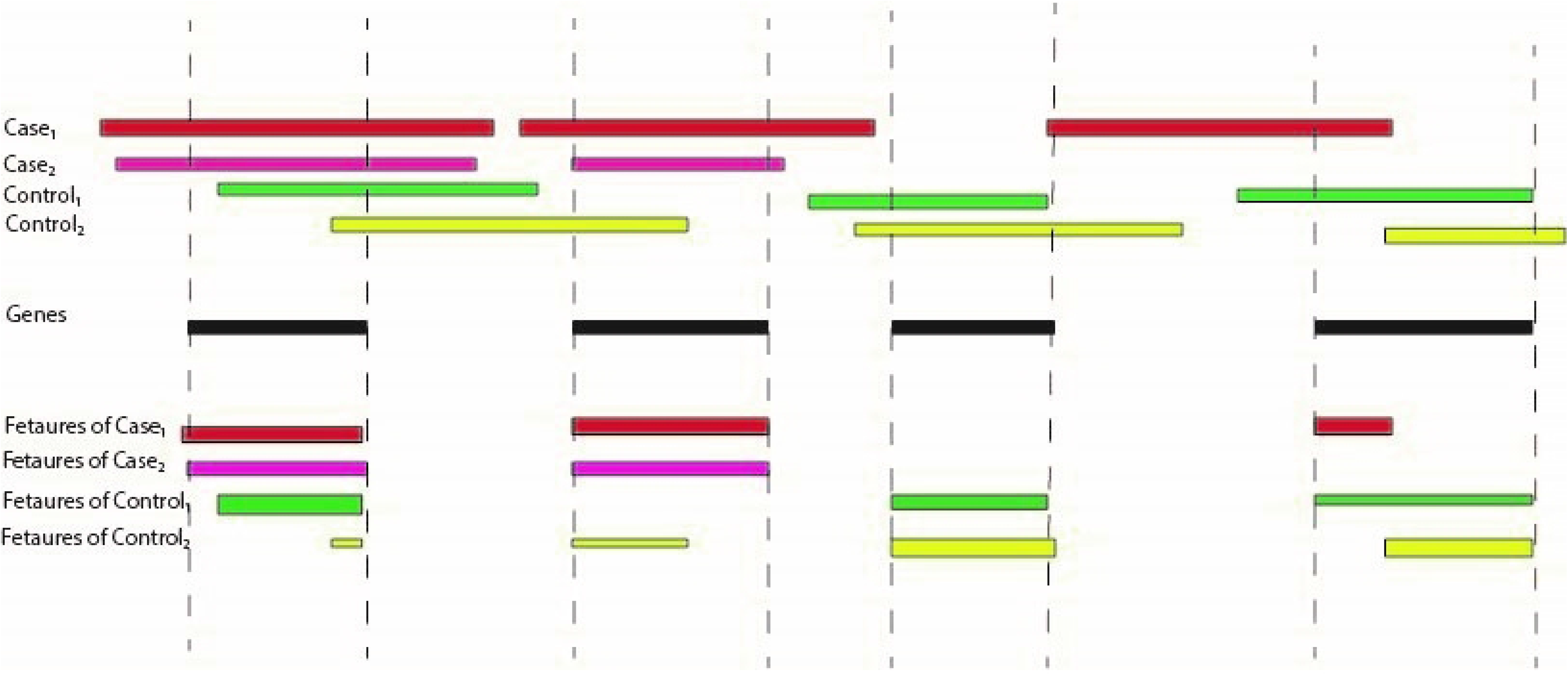
Heatmap for autism.

**Figure 7:**
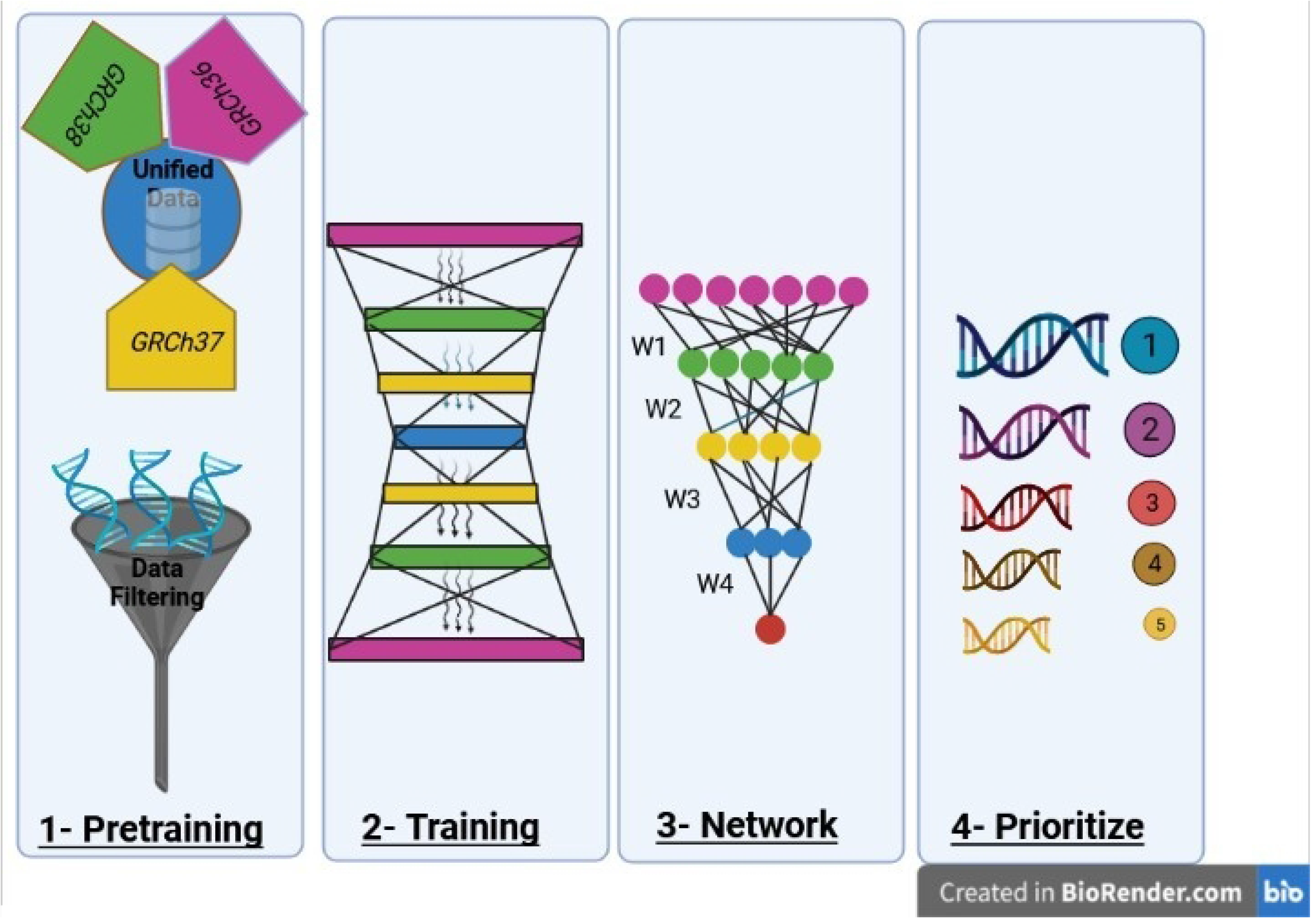
Distribution of CNV length in different chromosomes for SCZ disease; y − Axis is the ×10^5^. The numbers in the table on top of the plot show the number of cases and controls. The red color (left) represents cases, and the blue (right) represents controls.

**Figure 8:**
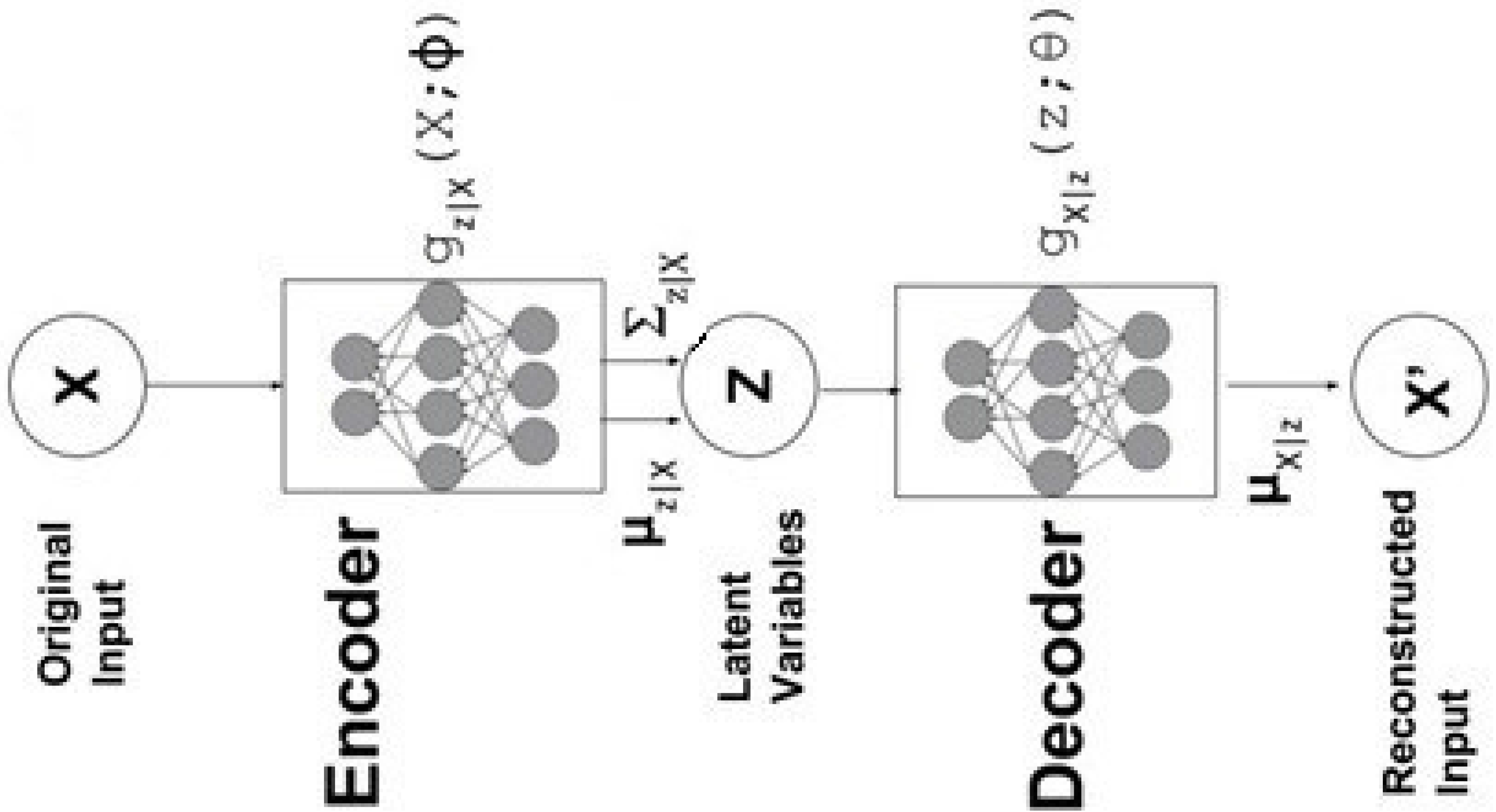
Distribution of CNV length in different chromosomes for ASD disease. Y-Axis is the ×10^6^. The numbers in the table on top of the plot show the number of cases and controls. The red color (left) represents cases, and the blue (right) represents controls.

**Figure 9:**
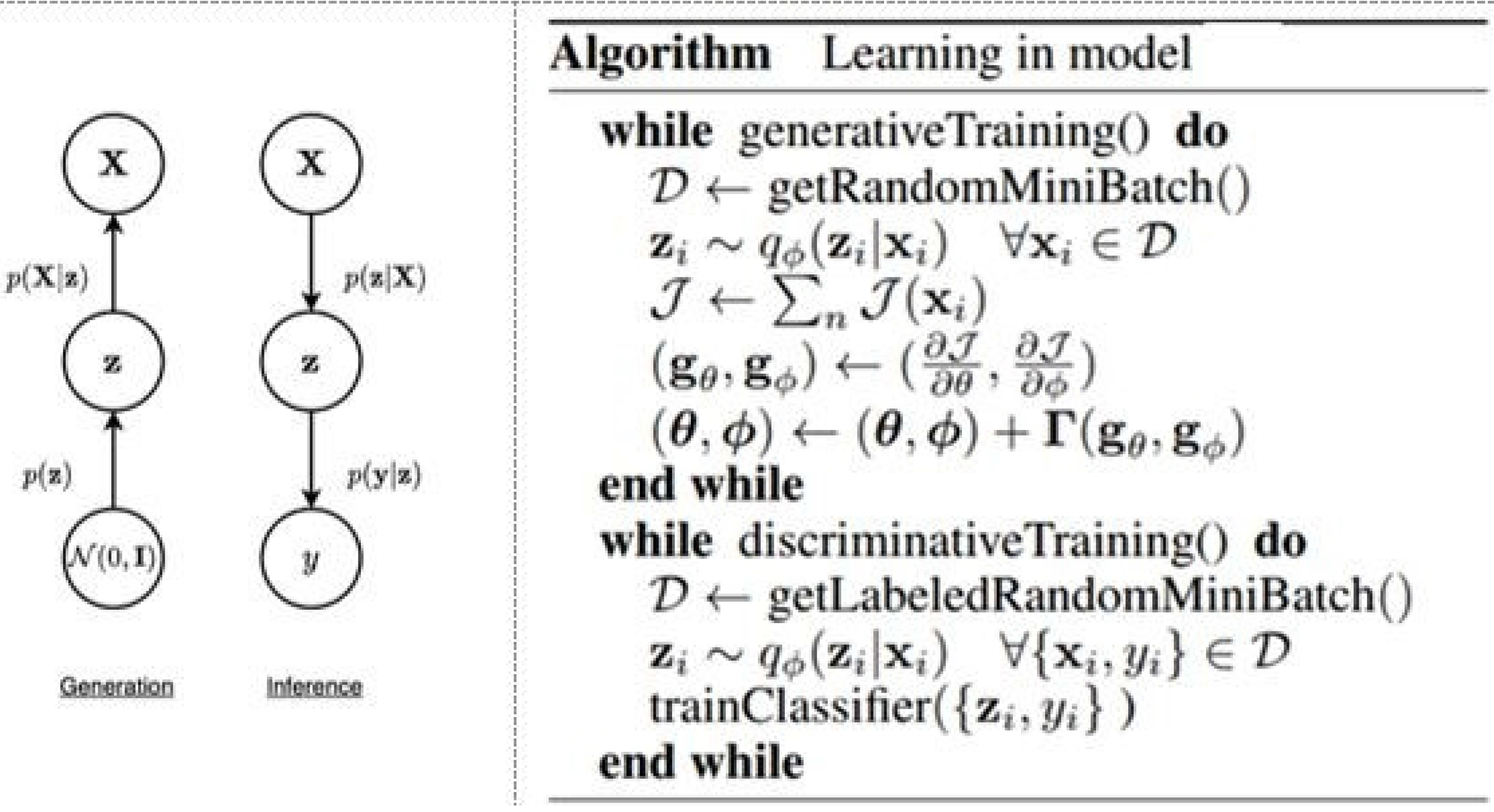
Distribution of CNV length in different chromosomes for DD disease. Y-Axis is the ×10^6^. The numbers in the table on top of the plot show the number of cases and controls. The red color (left) represents cases, and the blue (right) represents controls.

**Figure 10:**
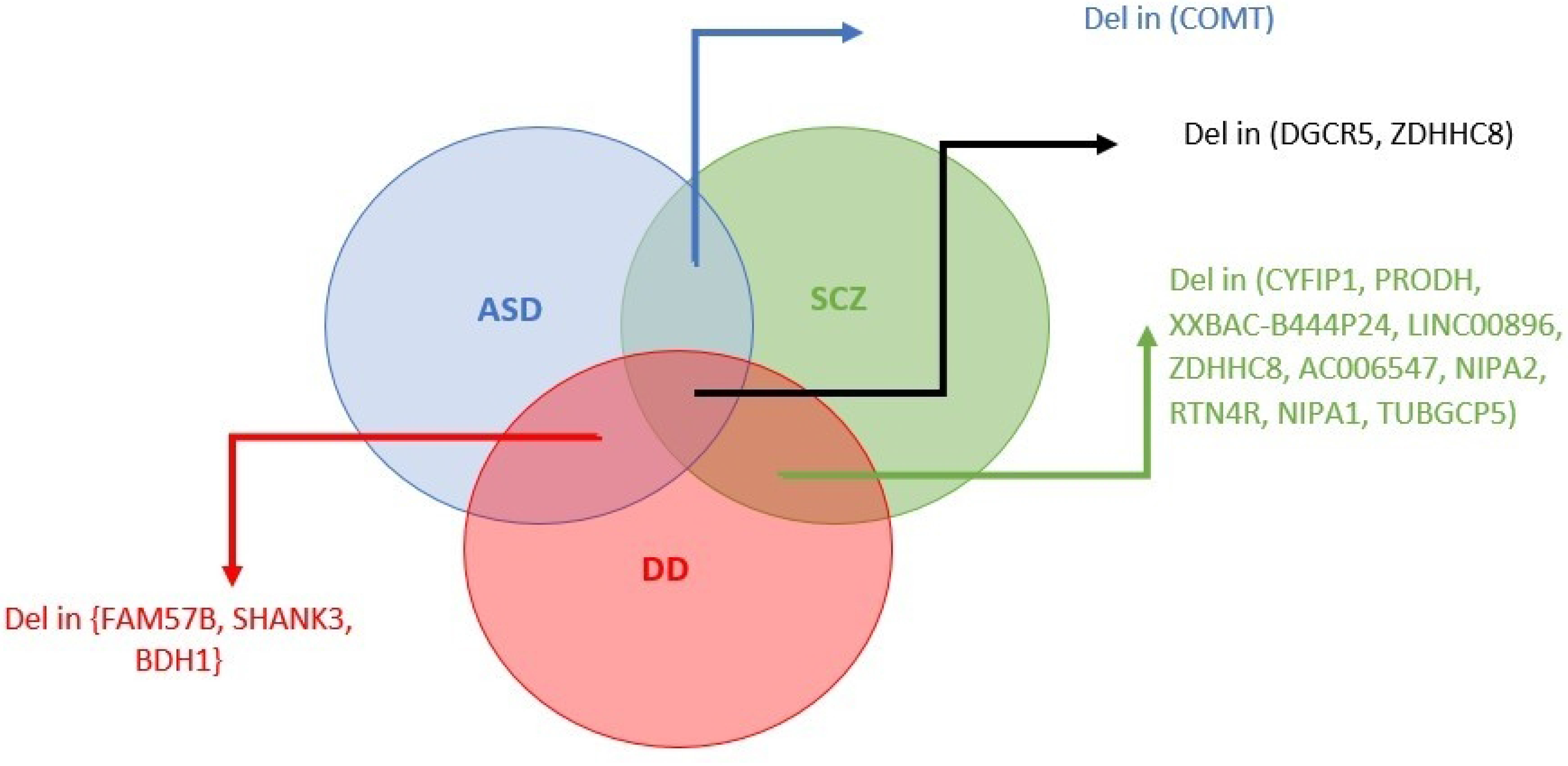
Visualization of a two-step semisupervised variational autoencoder(VAE) process; first, the VAE is trained in an unsupervised manner. Then one part is used to train with labels.

We also searched for genes with brain-enriched expression. Numerous genes with brain-enriched functions are associated with neurodevelopmental disorders [39]. Our findings were compared to those of similar works. Results from our study are more enriched than those from others.

Additionally, we compared the results to those genes that cause nervous system phenotypes in mice. Data were obtained from the MGI repository. Compared to similar studies, ours has a higher fold enrichment. Our next step is to identify genotype-phenotype relationships using the DECIPHER data source. Phenotypes with high enrichment levels are discovered.

Additionally, we utilized WebGestalt [35] to perform gene ontology analyses within coding genes to examine Gene Ontology (GO), Human Phenotype Ontology (HPO), or associated disease terms. Each biological observation is explained in detail in the following sections.

### Overrepresentation of Tissue-specific Genes

Several studies (such as [28] and [31]) claim that brain-enriched genes play an important role in NDDs. To determine whether the detected genes are overrepresented in the brain tissue, we measure the fraction of coding and non-coding genes that have been enriched (background expectation) and compare it with the percentage of genes that have overlapped with deleted or duplicated CNVs.

A list of brain-enriched genes is provided in [28]. For obtaining this list, they used the FANTOM5 CAGE-associated transcriptome [32] to identify coding and long non-coding RNA genes present in the regions and examined their expression patterns across sample types.

Table 2 presents the results of brain-enriched coding genes fold enrichment, while Table 3 illustrates brain-enriched lncRNA genes fold enrichment.

The results of our study are compared with those of Coe et al. [11] and Cooper et al. [33], two significant studies of developmental delay. Their gene expression is less brain-enriched than ours. The results were also compared with those of PLINK [34]. The next comparison tool is SNATCNV [28], a publicly available tool with state-of-the-art performance.

In the list of brain-enriched genes related to ASD and SCZ, *DGCR2* specifies a protein proposed to play a role in neural crest cell migration. The *ZDHHC8* gene, strongly associated with ASD and SCZ, is another gene to note.

Next, we have some brain-enriched genes associated with SCZ; *RTN4R* is a gene in which adult central nervous systems are likely to be affected by its role in regulating axonal regeneration and plasticity. *CATG00000058203* and *Septin5*, and *CATG00000057131* are some brain-enriched genes associated with ASD and SCZ, previously mentioned in [28].

As for the developmental delay, the *DGCR5, PRODH, NIPA1, TUBGCP5, RTN4R, ZDHHC8, CRKL,* and *SERPIND1* genes are also brain-enriched and associated with the disease. Most of them are from the 22^nd^ and 15^th^ chromosomes (22q11.21).

### Gene Segregation Analysis of Male and Female Patients

Long-standing research shows that females are more tolerant of mutations than males, which explains why males are more prone to neurodevelopmental disorders such as autism. New studies also confirm the validity of previous findings [53–55] that male cases show more significant enrichment than female cases when comparing the ratios of cases to controls. In this research, we pointed out that some genes are more biased towards males, for example, deletion in *PHF2 (ENSG00000197724)*, duplication in *NRXN1 (ENSG00000179915)*, and deletion in *WDFY3 (ENSG00000163625)*, *PHF3 (ENSG00000118482)*, *MED13L (ENSG00000123066)*, and *WAC (ENSG00000095787)*, are more frequently seen in males than females for the developmental delay.

Besides, we performed the same analysis with ASD CNVs. We found that the *PTCHD1 (ENSG00000165186)* gene deletion occurred more in male than female patients.

### Phenotype Association with Causal Genes of target Neurodevelopmental Disorders

To analyze phenotypes associated with causal genes, we used DECIPHER [27], the genotype-phenotype data source for almost 12,600 patients with CNVs; Fig. 5 represents the distribution of phenotypes in DECIPHER.

This data source has 1,548 patients related to ASD, with ‘HP:0000717’ (autism), ‘HP:0000729’ (autistic behavior), and ‘HP:0000753’ (autism with high cognitive abilities) phenotypes, and 2,144 patients related to DD with ‘HP:0001263’ (global developmental delay), ‘HP:0011342’ (mild global developmental delay), ‘HP:0011344’ (severe global developmental delay), ‘HP:0011343’ (moderate global developmental delay), and ‘HP:0012758’ (neurodevelopmental delay) phenotypes.

To investigate the relationship between genes and phenotypes, we calculate the ratio of overlapped samples with the specific phenotype to the number of overlapped samples for a putative gene. Figs 11 (DD), 12 (SCZ), and 13 (ASD) depict the respective heatmaps for each target disorder.

**Figure 11:**
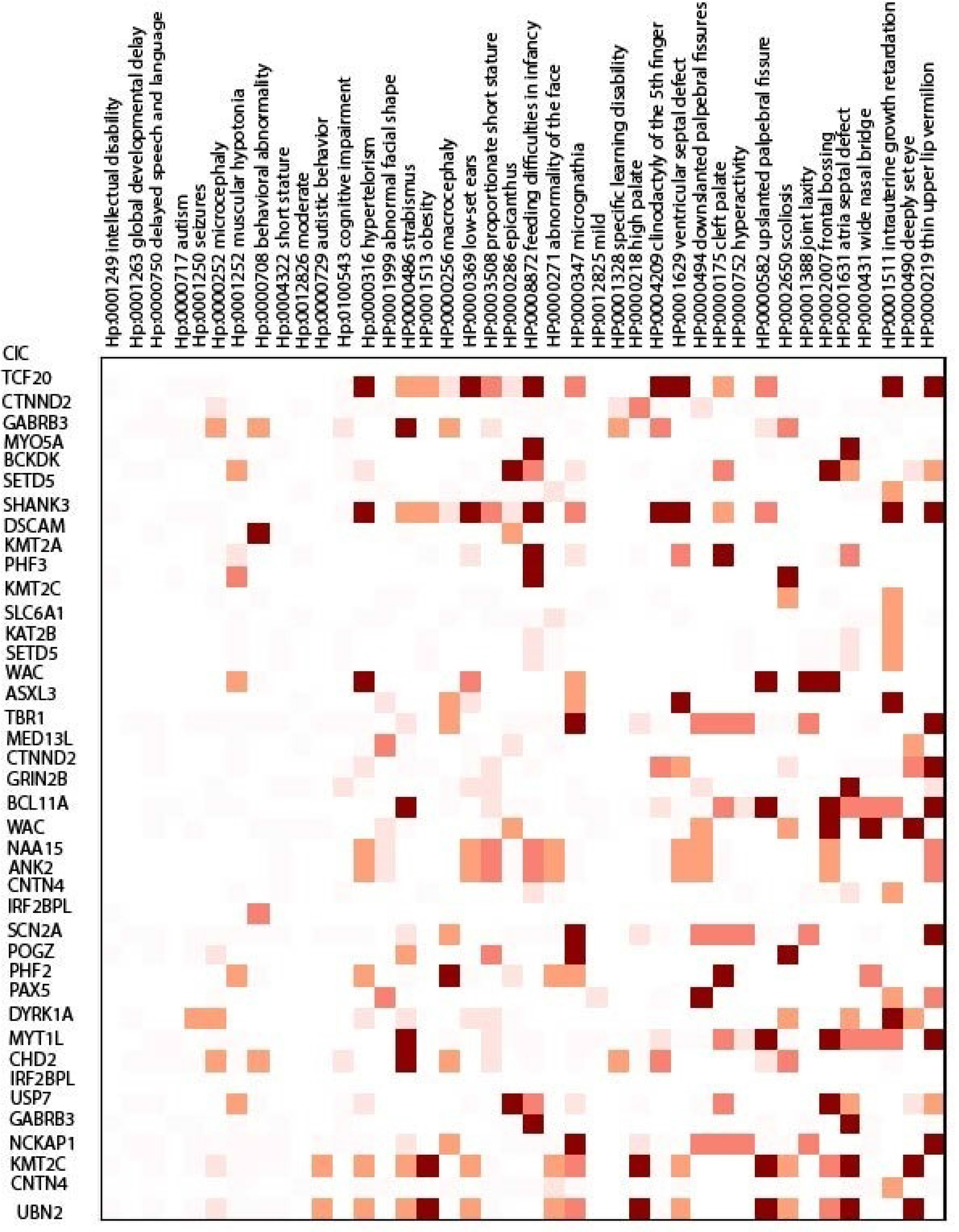
A sketch of the whole process; here, we tried to provide an overview of the proposed method. The first step (from the left) is a preprocessing step in data preparation. In this step, we have data in a different format (HG18, HG19, etc.), converting it to a unified one (HG19). We should remove some of the data that are redundant, useless, or include incomplete features. We should build a model out of the cleaned data in the second step. This model is a type of autoencoder, and in the second step, we edit the weights of the network with the labels (the labels are zero or one for the healthy and patient people). In the fourth step, we use the coefficients of the autoencoder for prioritizing the genes (the bigger icons represent the more important genes, and the smaller icons are representations of the less important ones).

**Figure 12:**
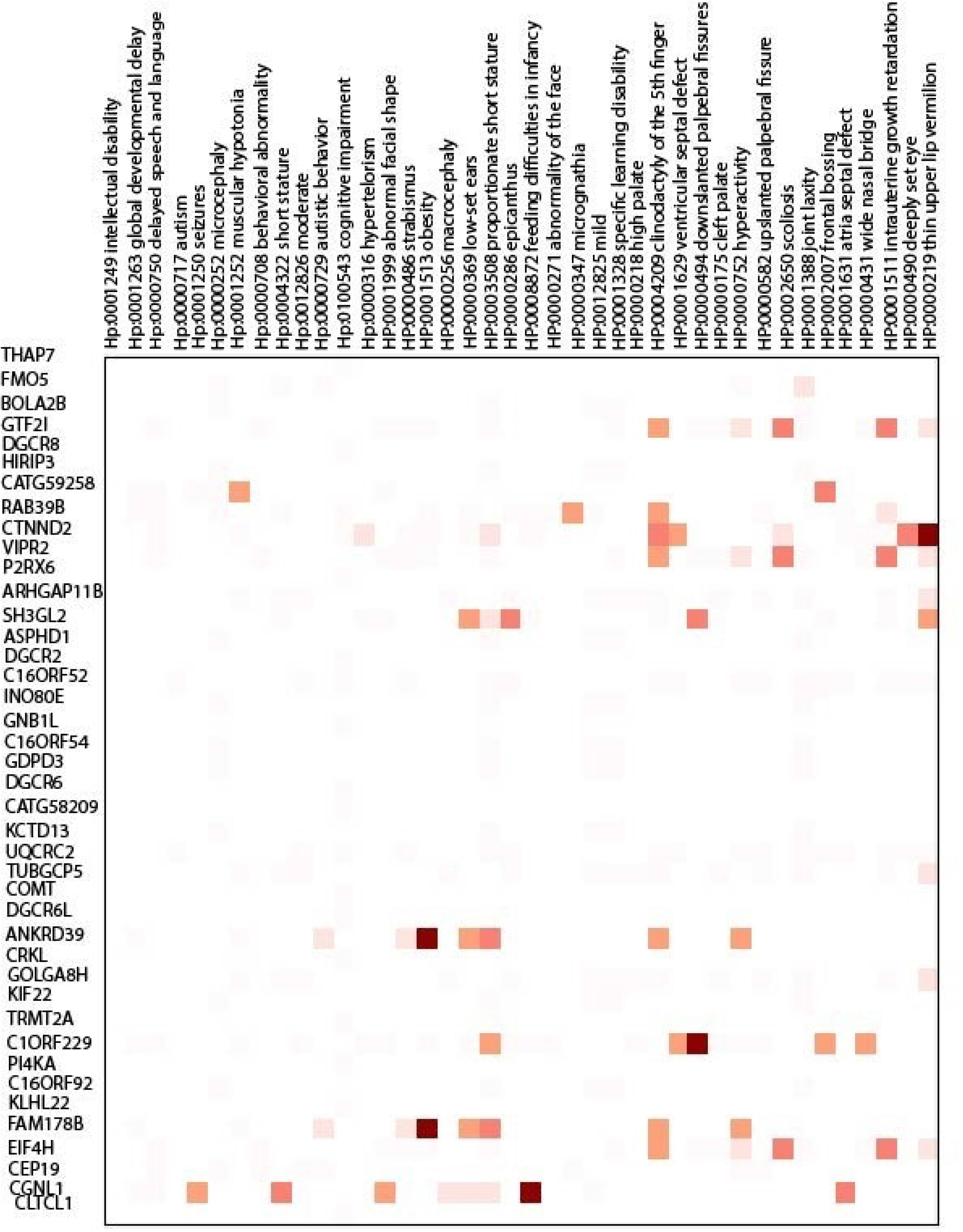
Features extracted from cases and controls; we have two cases and two controls in this figure and four genes. The CNVs of patients and healthy people are shown as rectangles, representing the starts and ends in a typical chromosome. The overlaps are shown at the bottom, which are features’ values for each case and control.

**Figure 13:**
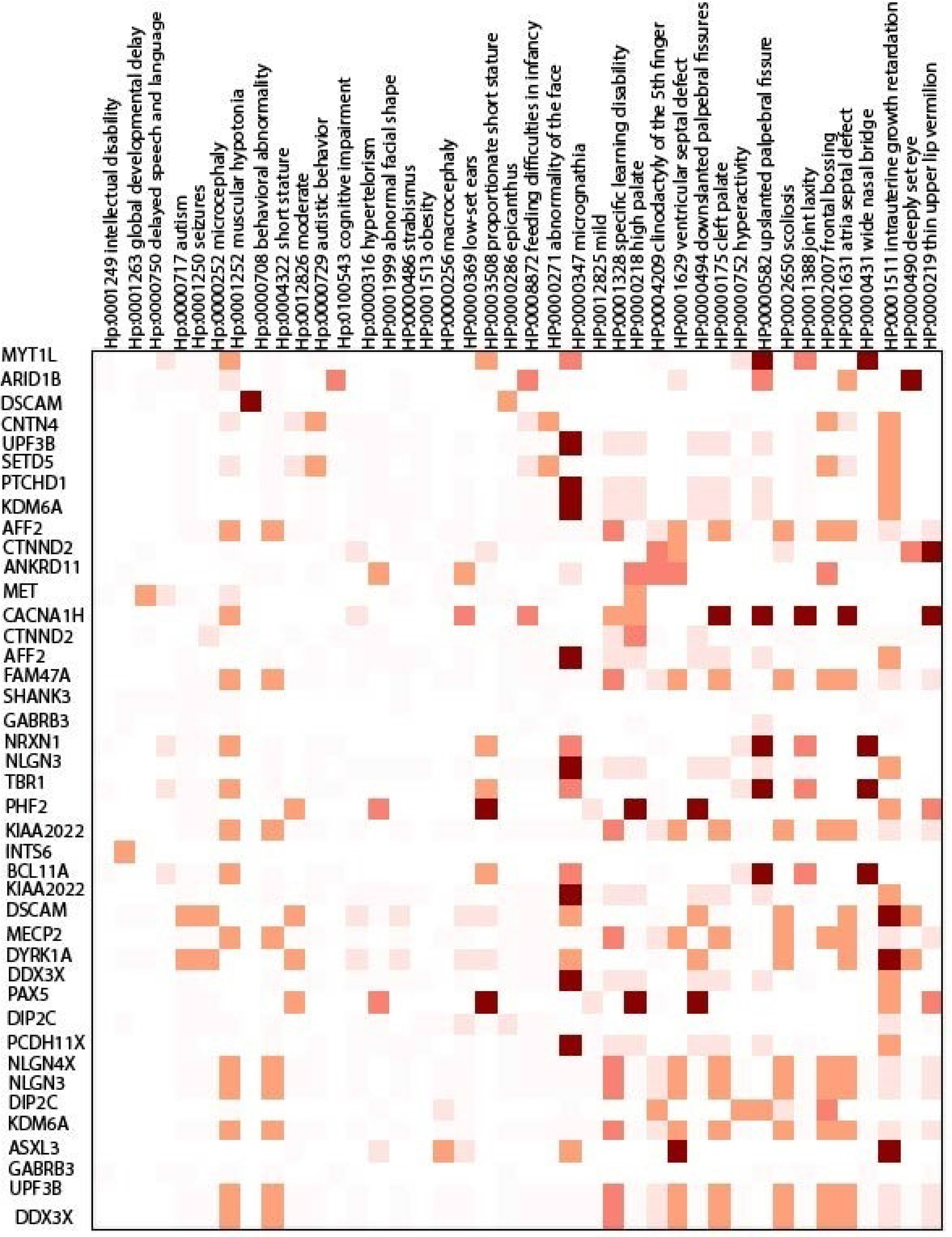
Variational autoencoder(VAE) (Left) A graphical model that clarifies the generation and inference process, (Right) The learning process algorithm [40].

Some of the highlighted phenotypes related to the target diseases are obesity (HP:0001513), autism (HP: 0000717), behavioral abnormality (HP: 0000708), irregularity of the face (HP: 00000271), and seizures (HP:0001250).

Children with autism are more likely to suffer from medical comorbidities. For example, we found macrocephaly (HP:00000256), hydrocephalus (HP:00000238), cerebral palsy (HP:0100021), migraine (HP:0002076), sleep disturbance (HP:0002360), and failure to thrive (HP:0001508) which was also mentioned in [56] as the phenotypes that co-occur in the autism.

For schizophrenia, DECIPHER analysis revealed phenotypes such as obsessive-compulsive behavior (HP:0000722), anxiety (HP:0000739), and depression (HP:0000716), as well explained in [57]. *MVP* duplication, overrepresented in SCZ, is associated with depression (HP:0000716).

Regarding the developmental delay, secondary conditions such as microcephaly (HP:0000252) and anxiety (HP:0000739) can be proposed, which was also suggested in [72]; This disorder has received less research. *BCL9*, *FMO5,* and *GPR89B* deletions related to microcephaly are also overrepresented in DD. *NIPA1* duplication, associated with anxiety, is among the top genes of DD. In [72], microdeletion of the *NF1* gene is found to be associated with microcephaly and DD.

Taken together, we have a model that deduces a set of genes for a target genetic disease. We investigated the set of phenotypes related to the genes; the particular relationship between genes and phenotypes shows that there can be diversity in the etiology of the disease, which implies that the occurrence of a phenotype in a target disease is influenced by what causal genes are mutated in the patient.

### Analysis of Biological Processes and Phenotypic Ontologies of Causal Genes

As part of our analysis, we used WebGestalt [35] to investigate the associations between identified genes and specific gene ontologies (GOs), human phenotype ontologies (HPOs), and disease terms [36], [37].

Some examples of the discovered disease ontology terms were autism spectrum disorders, intellectual disability, language development disorders, poor school performance (for the developmental delay), autistic disorder, and language development disorders. Language development disorders are discussed in [58] as a comorbidity of NDDs.

In the associated HPO terms, some examples were autistic behavior, delayed speech and language development, intellectual disability, severe global developmental delay, abnormal social behavior, impaired social interactions, and abnormally aggressive, impulsive, or violent behavior. Abnormal behavior is mentioned in [59], and impaired social interaction is discussed in [58] as phenotypes related to NDDs.

The highlighted Gene Ontology terms include cognition, dendrite development, and synapse organization. In [60], dendrite development is pointed out to be associated with NDDs, and the relationship between synapse organization and NDDs is addressed in [61]. Tables 12 through 17 summarize the results.

**Table 8:**
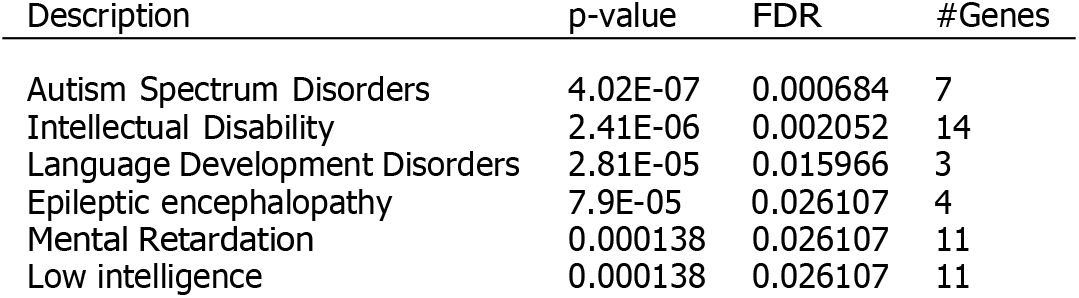
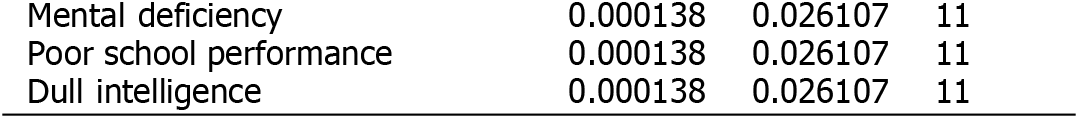
Disease Ontology Terms for developmental delay.

| Description | p-value | FDR | #Genes |
| --- | --- | --- | --- |
| Autism Spectrum Disorders | 4.02E-07 | 0.000684 | 7 |
| Intellectual Disability | 2.41E-06 | 0.002052 | 14 |
| Language Development Disorders | 2.81E-05 | 0.015966 | 3 |
| Epileptic encephalopathy | 7.9E-05 | 0.026107 | 4 |
| Mental Retardation | 0.000138 | 0.026107 | 11 |
| Low intelligence | 0.000138 | 0.026107 | 11 |
| Mental deficiency | 0.000138 | 0.026107 | 11 |
| Poor school performance | 0.000138 | 0.026107 | 11 |
| Dull intelligence | 0.000138 | 0.026107 | 11 |

**Table 9:**
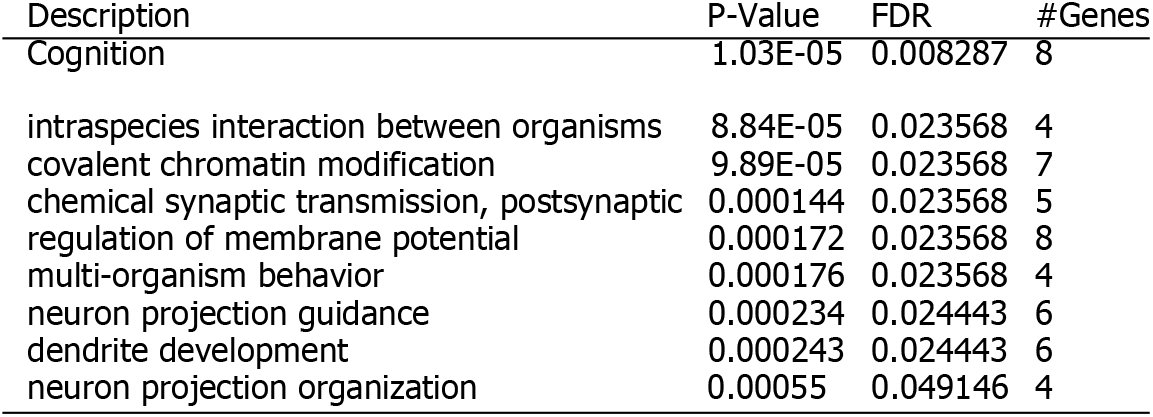
Gene Ontology for developmental delay.

**Table 10:** Disease ontology terms for schizophrenia.

| Description | P-Value | FDR | #Genes |
| --- | --- | --- | --- |
| Blepharophimosis | 4.04E-07 | 0.000429 | 5 |
| Hernia, Inguinal | 7.50E-07 | 0.000429 | 5 |
| Chronic otitis media | 1.01E-06 | 0.000429 | 4 |
| ear infection chronic | 1.01E-06 | 0.000429 | 4 |
| Proteinuria | 8.52E-06 | 0.000968 | 5 |
| Redundant skin | 9.1E-06 | 0.000968 | 3 |
| Bunion | 9.1E-06 | 0.000968 | 3 |
| Hallux Valgus | 9.1E-06 | 0.000968 | 3 |
| Sloping shoulders | 9.1E-06 | 0.000968 | 3 |
| Congenital anomaly of neck | 9.1E-06 | 0.000968 | 3 |
| Deformity of neck | 9.1E-06 | 0.000968 | 3 |
| Malformation of the neck | 9.1E-06 | 0.000968 | 3 |
| Hypoplastic toenails | 9.1E-06 | 0.000968 | 3 |
| Phonophobia | 9.1E-06 | 0.000968 | 3 |
| Colon diverticulum anatomic structure | 9.1E-06 | 0.000968 | 3 |
| Diverticular disease of the colon | 9.1E-06 | 0.000968 | 3 |
| Pointed chin | 1.05E-05 | 0.001048 | 4 |
| Sacral dimples | 1.81E-05 | 0.001621 | 3 |
| Pulmonary Stenosis | 2.3E-05 | 0.001621 | 4 |
| Prominent lower lip | 2.89E-05 | 0.001621 | 4 |

**Table 11:**
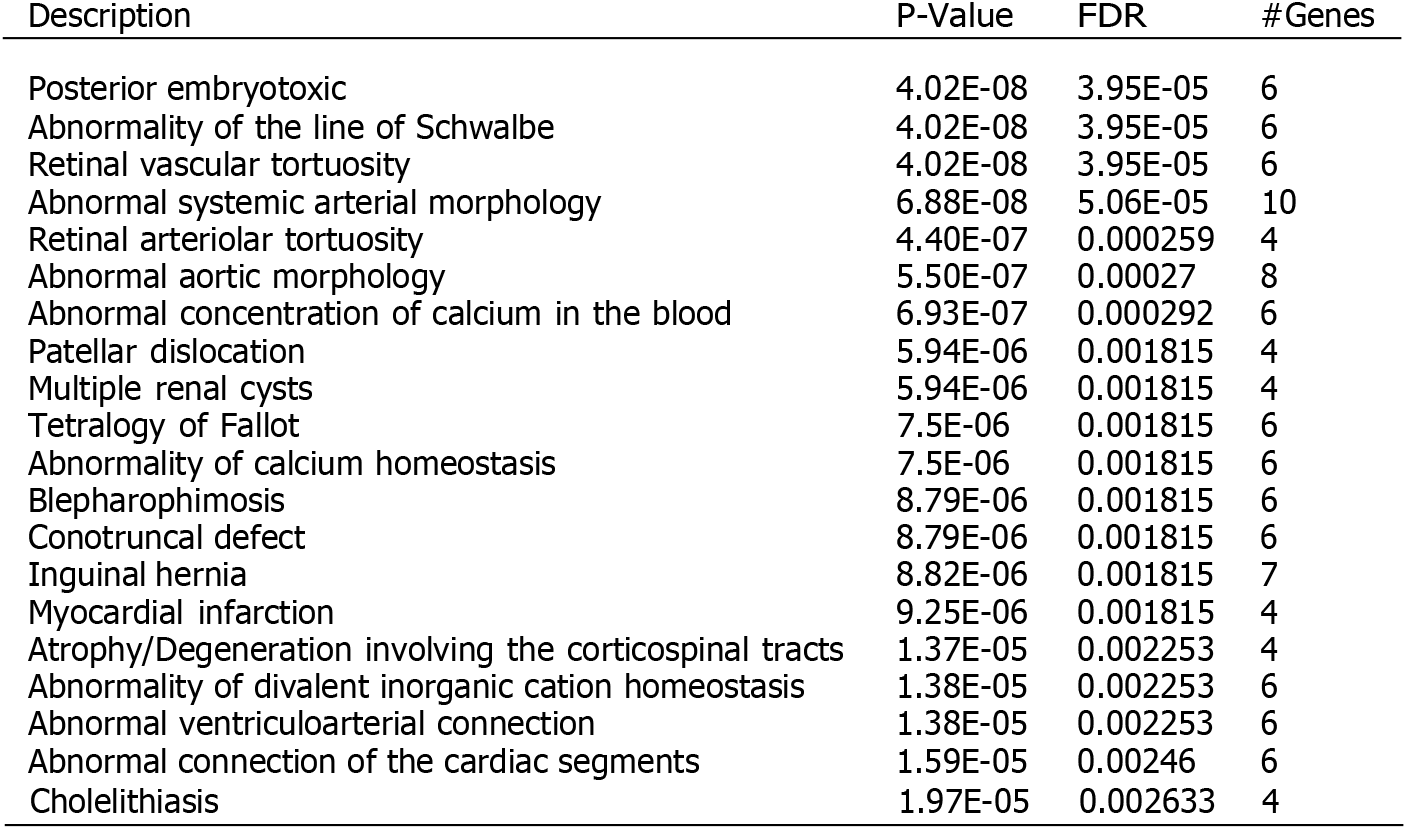
Human phenotype ontology terms for schizophrenia.

**Table 12:** Disease ontology terms for autism spectrum disorder.

| Description | P-value | FDR | #Genes |
| --- | --- | --- | --- |
| Autism Spectrum Disorders | 3.73E-08 | 6.3437E-05 | 7 |
| Autistic Disorder | 1.3592E-06 | 0.0011573 | 9 |
| Language Development Disorders | 0.000010744 | 0.006099 | 3 |
| Intellectual Disability | 0.000094389 | 0.038931 | 10 |
| Autistic behavior | 0.0001143 | 0.038931 | 3 |

**Table 13:** Gene ontology terms for autism spectrum disorder.

| Description | P-value | FDR | #Genes |
| --- | --- | --- | --- |
| Intraspecies Interaction Between Organisms | 8.79E-09 | 7.0694E-06 | 6 |
| Multi-Organism Behavior | 2.62E-08 | 1.0521E-05 | 6 |
| Cognition | 4.24E-07 | 0.00011364 | 8 |
| Chemical Synaptic Transmission, Postsynaptic | 9.33E-07 | 0.00018746 | 6 |
| Dendrite Development | 2.5364E-05 | 0.0040786 | 6 |
| Adult Behavior | 3.0537E-05 | 0.0040919 | 5 |
| Membrane Biogenesis | 6.3198E-05 | 0.0072205 | 3 |
| Synapse Organization | 7.1846E-05 | 0.0072205 | 7 |
| Regulation Of Membrane Potential | 8.3743E-05 | 0.0074811 | 7 |
| Respiratory Gaseous Exchange | 0.00022062 | 0.017053 | 3 |
| Peptidyl-Lysine Modification | 0.00023331 | 0.017053 | 5 |
| Neuron Projection Guidance | 0.00030854 | 0.020672 | 3 |
| Regulation Of Neurological System Process | 0.00048216 | 0.02982 | 4 |
| Covalent Chromatin Modification | 0.00085136 | 0.048892 | 5 |

**Table 14:** Mouse genes related to nervous system phenotypes.

|  | Deletion | Duplication |
| --- | --- | --- |
| Genome-wide Expectation | 12.7% | 12.7% |
| Our Result (DD) | 36.2% | 43.4% |
| Plink [34] | 12.0% | 13.7% |
| Our Result (SCZ) | 30.4% | 29.7% |
| SNATCNV (ASD)[28] | 14.4% | 14.4% |
| Our Result (ASD) | 31.3% | 40.5% |

**Table 15:** Statistics of different datasets.

\* 13k Affy and 15k Illumina for cases, 14K Affy, and 14k Illumina for controls.
| Dataset | of case CNVs | of control CNVs | Sum | of Patients | of Healthy | Sum |
| --- | --- | --- | --- | --- | --- | --- |
| Autism spectrum disorder | 47,119 | 24,858 | 71,977 | 19,663 | 6,479 | 26,142 |
| Schizophrenia | 42,046 | 40,414 | 82,460 | 28,684* | 28,893* | 57,577 |
| Developmental delay (NSTD 100) | 29,803 | 11,256 | 41,059 | 29,085 | 19,584 | 48,669 |

**Table 16:**
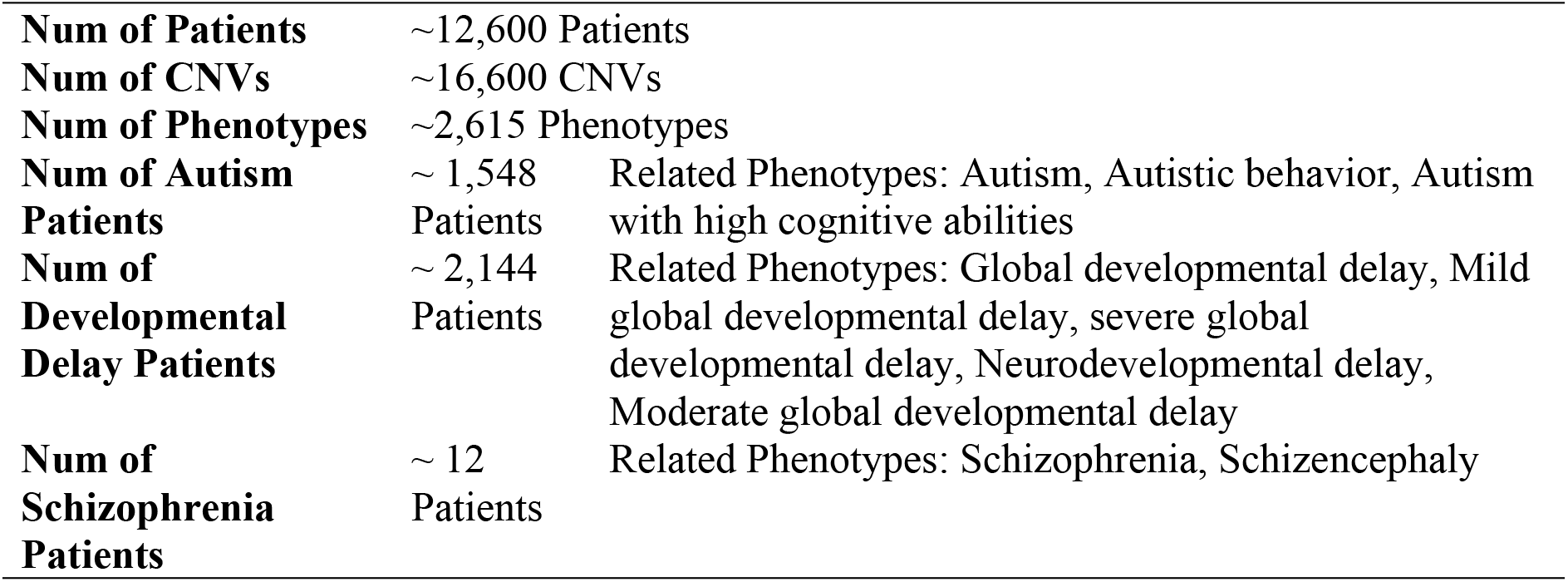
DECIPHER statistics [27].

|  |  |  |
| --- | --- | --- |
| **Num of Patients** | ~12,600 Patients |  |
| **Num of CNVs** | ~16,600 CNVs |  |
| **Num of Phenotypes** | ~2,615 Phenotypes |  |
| **Num of Autism Patients** | ~ 1,548 Patients | Related Phenotypes: Autism, Autistic behavior, Autism with high cognitive abilities |
| **Num of Developmental Delay Patients** | ~ 2,144 Patients | Related Phenotypes: Global developmental delay, Mild global developmental delay, severe global developmental delay, Neurodevelopmental delay, Moderate global developmental delay |
| **Num of Schizophrenia Patients** | ~ 12 Patients | Related Phenotypes: Schizophrenia, Schizencephaly |

**Table 17:** Supplementary data sources statistics.

|  | Coding Genes | lncRNA Genes |
| --- | --- | --- |
| Fantom5 Genes | 21,069 | 27,920 |
| Brain-Enriched Genes | 7,339 | 7,167 |
| MGI Genes | 3,427 | 0 |

### Overrepresentation of homologs of coding genes causative of nervous system phenotypes in the mutated mouse

Studying animal genetic mutations provides insight into disease mechanisms and treatments for neurodevelopmental disorders. Several animal models have been developed to uncover the disorder’s process [38]. Mutant mice with specific defects in the nervous system are among them. Models based on mutant mice replicate key symptoms of neurodevelopmental disorders.

We investigate what percentage of the causative genes have homologs in mouse genes whose mutation causes nervous system phenotypes. For this purpose, we used the Mouse Genome Informatics (MGI) database to identify genes related to the mouse nervous system and their human homologs.

Table 4 examines the fractions of homologs of the discovered genes that manifest nervous system phenotypes in mice. The results show that the coding genes detected by our method, compared to the results of other methods, have a higher percentage of homologs in mutant mouse models with nervous system phenotypes. For example, some of the genes that have orthologs in mice with nervous system phenotypes are *SEPTIN5, RTN4R,* and *ZDHHC8*.

These genes have high ranks and are common among the three disorders.

## Discussion

We present a method that utilizes deep learning to systematically analyze CNV datasets associated with a target disease (e.g., a neurodevelopmental disorder) and prioritize genes. The deep learning model learns how features are distributed over samples using an autoencoder architecture with a variational learning framework. Using this method, we can predict the likelihood that a gene mutation will cause a specific disease. We applied the technique to three neurodevelopmental disorders and examined the results to find their overrepresentation of enriched brain coding, long non-coding RNA genes, and mouse orthologs with nervous system phenotypes. Additionally, with DECIPHER data source, we investigated how mutations in the detected genes influence other traits, and gene ontology analyses were also conducted.

We had 118,968 case CNVs for 48,748 patients and 76,528 control CNVs for 26,063 healthy people for the CNV association with neurodevelopmental disorders.

In the top 40 genes for developmental delay, *DGCR6* is a candidate for involvement in DiGeorge syndrome pathology and schizophrenia; defects in *PRODH* (Proline Dehydrogenase 1), located on 22q11.21, are associated with susceptibility to schizophrenia 4 (SCZD4). *DGCR5* (DiGeorge Syndrome Critical Region Gene 5), an RNA gene associated with DiGeorge Syndrome; and expression of *MVP* (Major Vault Protein) may be a prognostic marker for several types of cancer. Additionally, defects in *ZDHHC8* may contribute to schizophrenia.

For schizophrenia, *DGCR6*, a protein-coding gene, is a well-known candidate for this disease. Mutations in *PRODH* (Proline Dehydrogenase 1), located on 22q11.21, are linked with susceptibility to schizophrenia 4 (SCZD4). *DGCR5* is a lncRNA that has a high score for causing schizophrenia. *SEZ6L2* (Seizure Related 6 Homolog Like 2), located in 16p11.2, where the region is thought to hold candidate genes for autism spectrum disorder. *CDIPTOSP* (CDIP Transferase Opposite Strand, Pseudogene) is a lncRNA gene associated with Central Nervous System Germ Cell Tumor disease. *ASPHD1* (Aspartate Beta-Hydroxylase Domain Containing 1) associated with Schizophrenia 3. *RANBP1* (RAN Binding Protein 1), a Protein Coding gene, is linked with Digeorge Syndrome.

For the autism spectrum disorder, *DGCR2* deletion is associated with a wide range of developmental defects (like DiGeorge syndrome, velocardiofacial syndrome, conotruncal anomaly face syndrome, and isolated conotruncal cardiac defects) categorized with the acronym CATCH 22; The *ARVCF* gene is responsible for the autosomal dominant Velo-Cardio-Facial syndrome (VCFS), with phenotypic features including cleft palate, conotruncal heart defects, and facial dysmorphology; *GNB1L* is also deleted in DiGeorge syndrome; *COMT* (Catechol-O-Methyltransferase), a protein-coding gene, associated with schizophrenia and Panic Disorder 1; *ZDHHC8* (Zinc Finger DHHC-Type Palmitoyltransferase 8), a protein-coding gene is linked to schizophrenia and Chromosome 6Q24-Q25 Deletion Syndrome; epilepsy is thought to be linked with *CHRNA7*, and mutations in *NRXN1* are connected with Pitt-Hopkins-like syndrome-2 and may increase the risk of schizophrenia.

We investigate which genes were more overrepresented in one gender and less in another. Duplication in *NRXN1 (ENSG00000179915)* and *PTCHD1 (ENSG00000165186)* gene deletion are more frequently seen in males than females for some of the NDDs.

We reported some brain-enriched coding genes were significantly expressed in all three disorders. Some examples were *DGCR2, SEPTIN5,* and *ARVCF* in the 22nd chromosome with the deletion. These three were among the top ten coding brain-enriched genes associated with three disorders. *DGCR5*, a noncoding brain-enriched gene (previously known as the biomarker of Huntington’s disease [62]), is proposed as highly associated with DD. *AC000068* is the noncoding brain-enriched gene associated with SCZ and ASD. Previously, *SEPTIN5* was shown to be associated with ASD and SCZ [63]. *DGCR2* was mainly known to be associated with SCZ [64]. *AC004471* is the noncoding brain gene among the top 10 genes related to SCZ, ASD, and DD.

Among the top genes with significant brain expression, some have orthologues in mice that showed nervous system phenotypes. *SEPTIN5, ZDHHC8, RTN4R,* and *KCTD13* are the four top genes for ASD and SCZ; *RTN4R* and *ZDHHC8* also rank highly in DD. *SEZ6L2* is top in ASD but with a lower rank in SCZ. *ZDHHC8* and *RTN4R* are genes with the nervous system, morphological, and physiological phenotypes. *SEPTIN5* shows only nervous and physiological phenotypes in mice.

In the next step, DECIPHER was used to examine the relationship between detected genes and other phenotypes; delayed speech, language, and autism were associated with the genes. According to our findings, seizures were associated with SCZ; this relationship was previously discussed in [65].

Microcephaly [66] and macrocephaly [67] (the two reverse phenotypes) are associated with ASD and SCZ. ‘Abnormal facial shape’ is associated with all three disorders [68]; it is also studied in [69]. *CACNA1H* is one of the genes related to some overrepresented phenotypes [70]. *TCF20*, discussed in [71], is another gene highlighted in the heatmap of developmental delay.

Gene ontology analysis for the detected genes was performed with the WebGestelat tool. With the help of this tool, gene ontology analysis, human phenotype ontology (HPO) analysis, and disease ontology analysis were performed separately. For the disease ontology, some terms were ‘Language Development Disorders,’ ‘Autistic behavior,’ and ‘Congenital anomaly of the neck.’ Overrepresented human phenotype ontology terms included ‘Severe global developmental delay,’ ‘abnormal social behavior,’ ‘Delayed speech and language development,’ and ‘Intellectual disability.’ And of the most common gene ontology terms were ‘dendrite development,’ ‘cognition,’ ‘Regulation of Neurological System Process.’ In summary, these findings support the biological relevance of the tool-identified CNV regions to genetic factors that contribute to neurodevelopmental disorders.

## Conclusions

To explore the effect of genes in neurodevelopmental disorders, we developed a tool based on deep learning for analyzing genes responsible for a target disease. We trained our model with all of the CNVs from the three neurodevelopmental disorders; thus, we made the most effective use of data in the pretraining phase; after that, we used CNVs of the target disease in the next stage for fine-tuning. We compared the results with some of the related works for each of the target diseases. The discovered genes include more coding and lncRNA ones enriched in the brain, and our results have more homologs in the mouse with nervous system phenotypes. Besides, we used the DECIPHER data source to identify the phenotypes related to the genes of the target disease. Integration with the phenotypic database revealed more attractive characteristics of the detected genes.

In the future, we can model gene relationships with graph-based models and do classifications. An alternative future path is to use additional evidence, such as protein networks to analyze the association of CNVs with diseases. Additionally, DECIPHER data, a multi-phenotype data source with CNVs for each patient, can provide a basis for analyzing the relation of the genetic etiology of the disease with the observed phenotypes in the patient and the possible co-occurrence of some phenotypes. Based on solid theoretical foundations, our method uses CNV data and can identify genes related to a specific disease.

## Materials and Methods

### Data and Preprocessing

We analyzed three case-control studies of neurodevelopmental disorders; autism spectrum disorder, schizophrenia, and developmental delay. Autism spectrum disorder data contains 47,119 case and 24,858 control CNVs documented in AUTDB database [28]; Schizophrenia consists of 42,046 case and 40,414 control CNVs [29], and Developmental delay consists of 29,803 case and 11,256 control CNVs (distribution of the CNVs length for the SCZ, ASD and DD are shown in Figs. 2, 3 and 4)[11].

The last data source, the developmental delay, has two data types in two independent datasets, NSTD 54 [33] and NSTD 100 [11]. NSTD 54 is a multi-phenotype data source that contains CNVs of ‘abnormality of the nervous system’ and ‘abnormal facial shape’ and ‘seizures’ and ‘generalized seizures,’ ‘autism,’ ‘failure to thrive,’ ‘abnormality of the cardiovascular system,’ ‘attention deficit hyperactivity disorder,’ ‘hearing impairment,’ ‘abnormality of the kidney,’ with high frequency. NSTD 100 is a developmental delay data source with gender data. Here, we used NSTD 100. All data are rare CNVs (frequency found in less than 1% of the population). The details are reported in Table 5.

The other supplementary data we have used is FANTOM5 (Functional Annotation of the Mammalian Genome 5) Atlas [41], consisting of 21,069 coding and 27,920 non-coding genes.

The next data source is the Database of Chromosomal Imbalance and Phenotype in Humans Using ENSEMBL Resources (DECIPHER, February 1st, 2017) [27]. This data source contains data on patients, CNVs, and phenotypes, namely ASD, DD, and SCZ. The statistics are shown in Table 7. We have used this data to analyze the relation of genes and other phenotypes; besides, data can be used for augmentation and pretraining of the system.

In this paper, we also work with tissue-enriched genes; they are genes with a high expression level in a particular tissue compared to all other tissues. The specific tissue here is the brain. We used the list of brain-enriched genes provided in [28]. The effect of brain-enriched genes on autism spectrum disorder is emphasized in [31]; here, we want to investigate their impact on schizophrenia and developmental delay.

Furthermore, MGI (Mouse Genome Informatics) data [42] determines whether causal genes related to disease cause a nervous system phenotype in the mouse. This work is also performed in [28]. Similar to [28], we want to find out whether the mutation of a gene orthologue causes a mouse mutant phenotype or not. In [28], HTML was parsed from the pages covering, (i) nervous system phenotype (MP:0003631) [43], (ii) abnormal nervous system morphology (MP:0003632) [44], (iii) abnormal nervous system physiology (MP:0003633) [45]; and the mapping was performed with [46]. A summary of the data sources is provided in Table 6. Preprocessing of data can be categorized into two parts; CNV filtering and conversion and supplementary data cleansing (DECIPHER data analysis, FANTOM5 data …).

For the CNV filtering and conversion, CNVs smaller than 1kbps were filtered (this filtering is performed in similar works like [33, 11, 28]), and different CNV studies were in other coordinates (hg17, hg18, and hg19). All the CNVs were first unified and converted to hg19 using UCSC Lift Genome Annotations tools [47]. Besides, the Y chromosome CNVs were filtered because there was insufficient data, eliminating all CNVs with missing values.

In supplementary data cleansing, those patients without any phenotypes were removed for using the DECIPHER data. No special preprocessing was needed for the Fantom5, MGI, and brain genes; all gene coordinates are in HG19 format, ready for processing.

Additionally, some genes that were not the result of the model were removed. These include genes that overlap more with controls than cases or genes that do not overlap with CNVs.

### A Formal overview of a gene prioritization system

If we imagine gene prioritization as a system, the input is a target disease plus the list of all genes; depending on the nature of the method that is used to process the genes, different datasets may also be utilized as the auxiliary input; for example, mutation data (SNPs or structural variants), protein networks, pathway data, or reliable causal genes related to a target disease (to use with the principle of ‘guilt by association’). The output is the list of candidate genes (which may be sorted or unsorted, the result would be either prioritization or classification). Also, a score may indicate the likelihood of a gene responsible for a phenotype (or a disease). The discriminatory algorithm tries to infer each gene’s role in the target’s incidence.

Here, we want to present a formal definition of our work. Let’s suppose we have a case-control study related to a target disease. This study includes copy number variants (CNVs) of patients and healthy people. CNVs are quadruples defined as follows:

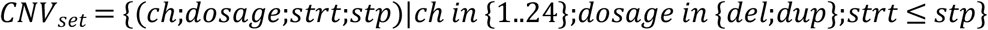

Where ch is the chromosome number, the dosage is the type of CNV, either deletion or duplication, and strt and stp determine the region of the chromosome where the variation occurs.

This CNV is available in two sets: one for cases and one for controls.

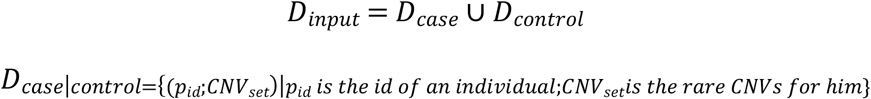

Each rare CNV is related to an individual (characterized by p_id), either healthy or patient. Optionally, the dataset may provide auxiliary data for the individual, such as gender information, which helps us investigate the discriminatory role of genes on each gender. Our goal is to solve the gene prioritization problem using the set of rare CNVs.

### The Method Overview

Compared to traditional machine learning methods, deep learning methods help build a pyramid of features and reduce data dimensions in a hierarchy. This will allow us to discover hidden patterns in data more effectively than other approaches. Autoencoder, a form of deep learning, reduces data dimension and produces a high-level, efficient view of the data from a hierarchy of features [6]. It has an encoder network (inference network) that gradually deconstructs input into a low dimensional latent representation and then a decoder network (generative network) that tries to reconstruct the output as similar as possible to the input. The autoencoders are used in many bioinformatics problems [48, 49, 50]. VAE [25]-[51], a combination of autoencoders and the variational learning framework, successfully extends the autoencoder. The training algorithm of the autoencoder is summarized in Fig. 8.

The main difference between autoencoders and variational version is that the first is deterministic, whereas the second is probabilistic; a variational autoencoder is an autoencoder whose training set is somehow regularized to avoid overfitting.

VAE is based on the Bayesian theorem and inference, with regularization constraint, assuming that latent representation has a multivariate Gaussian, N (µ, σ).

It is shown that VAE is more stable during training and has less vague output than other generative models; since it optimizes precise objective functions based on likelihood [52]. The posterior is a Gaussian distribution, whose output is mean and variance, and it is proved that all functions can be approximated with it. The model tries to encode the inputs into a Gaussian distribution with estimated mean and covariance.

VAE, a deep generative model based on variational inference, is a tool for finding a low-dimensional latent representation, z, of high-dimensional input data X, with the distribution P (X). To capture the inherent information of the input dataset, P(z|X), the posterior distribution is estimated, which is usually intractable. With the help of the variational inference, we have a distribution family Q(z|X) (variational distribution) for estimating the P (z|X) distribution. The approximation minimizes the Kullback–Leibler (KL) divergence (D) between the two distributions.

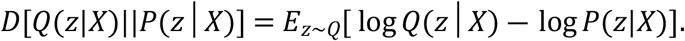

After some calculations, we have the following objective function, which is the variational lower bound on log-likelihood:

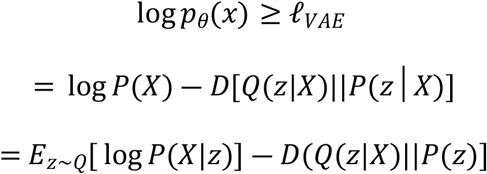

The first term is the expectation over the approximate posterior distribution (named as reconstruction error), and the second term (KL distance) is the regularizer (P (z) is standard Gaussian Distribution, N(0, I)). Q(z|X) is the encoding distribution, and P (X|z) is the decoding distribution.

With these equations, minimization is converted to maximization. Q(z|X) is the encoder, and P (X|z) is the decoder. The deep neural network can achieve this goal with stochastic gradient variational Bayes. The encoder part of VAE is used to produce the parameters of the variational distribution. The dropout technique can be applied to avoid overfitting. Q(z|X) (recognition model) is multi-dimensional Gaussian distribution, and the network will generate the mean and covariance of the Gaussian distribution. Standard Gaussian distribution N (0, I) is used as the prior for latent space.

The loss function comprises reconstruction loss (similar to autoencoder for efficient encoding-decoding action) and regularization term (also called latent loss, for restriction on latent space). The regularization term approximates the latent space as a standard Gaussian distribution. VAE adds Kulback-Leibler divergence to the loss function, requiring the covariance matrix close to identity and the mean to zero.

The training of the deep learning models consists of two phases: pretraining and fine-tuning. In pretraining, the autoencoder learns high-level features of all of the CNVs of the disorders. In fine-tuning, we put aside the decoder and use the dedicated CNVs for the target disease.

### The Method Details

In this section, we explain our method for prioritizing genes. An overview of the method is provided in Fig. 7.

A deep learning model is proposed for this task. According to the dataset for each disease, we have a copy number of variants for patients and healthy individuals. Each set of copy number variants for an individual has some overlaps with genes, and these overlaps are features that feed into our deep learning. This is shown in Fig. 6.

We have a list of genes that we want to determine whether their expression will affect disease incidence; on the other hand, we have a list of cases and controls with CNVs for a target disease. We want to convert them to a supervised learning algorithm.

In healthy and patient individuals, we want to convert CNVs to genes. Computing overlaps can do this. For the set of genes, preprocessed as discussed before, we measure the length of overlap (in kbps) with the CNVs of an individual. The label of the training set is whether the person is sick or healthy (zero or one).

In the pretraining phase of the model, we used all the CNVs of the neurodevelopmental disorders (autism + schizophrenia + developmental delay). The CNV of a specific disease is used for fine-tuning. This can be considered a form of semi-supervised learning.

After our VAE has been fully trained, we just use the encoder part directly for the next step:

1. Train a VAE using all our data points and transform our data (X) into the latent space (Z variables) (We use all data in this step).
2. Solve a standard supervised learning problem with (Z, Y) pairs (Y is the label set).

A graphical model and its learning algorithm for the whole process are shown in Fig. 9.

Let’s suppose that the encoder weights are represented by 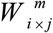, where m is the layer number, i is the output size in the last layer, and j is the input size in the current layer (no connection is determined by zero). As we know, the final layer that will be attached to the encoder is the label; and its size is one (whether the individual is patient (=one) or healthy (=zero)).If we multiply all weights matrices together, the result has the size input size × 1 (the matrices are multiplicable since the output of the last layer equals the input of the current layer). The resulting matrix (specifically column vector) can rank genes according to the label (the label is the status of the disease), and this is the same thing we want to model. The formulation is as follows:

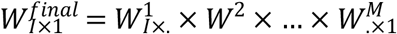

The specification of the deep learning model is such that a binary classification task is accomplished. The final layer has a binary outcome, the last activation function is sigmoid, and loss function is binary cross-entropy, and the optimization algorithm is Adam.

### The implementation Details

The deep learning model has a training phase, which needs a training set that includes the cases and controls. We developed the system with python and PyTorch [50]. We used cross-validation and grid search to tune the parameters, like the number of neurons in each layer. The activation functions are empirically-selected Rectified Linear Units, and the weights were optimized by an adaptive optimization algorithm (Adam) [43] to minimize reconstruction error and loss. The decoder has a symmetrical structure to the encoder. The learning rate, decay rate, and epoch were set to 0.001 and 1 and 10,000, respectively. Also, we restrict connections to some extent for a reduction in parameters. The train/test ratio is set to 80/20.

## Acknowledgments

Hamid R. Rabiee and Zahra Rahaie were supported by IR National Science Foundation (INSF), Grant No. 96006077.

## References

[1] Raj MR, Sreeja A. Analysis of computational gene prioritization approaches. Procedia computer science. 2018 Jan 1;143:395–410. https://doi.org/10.1016/j.procs.2018.10.411

[2] Lan W, Wang J, Li M, Peng W, Wu F. Computational approaches for prioritizing candidate disease genes based on PPI networks. Tsinghua Science and Technology. 2015 Oct 13;20(5):500–12. https://doi.org/10.1109/TST.2015.7297749

[3] Kumar AA, Van Laer L, Alaerts M, Ardeshirdavani A, Moreau Y, Laukens K, Loeys B, Vandeweyer G. pBRIT: gene prioritization by correlating functional and phenotypic annotations through integrative data fusion. Bioinformatics. 2018 Jul 1;34(13):2254–62. https://doi.org/10.1093/bioinformatics/bty079

[4] Nitsch D, Gonçalves JP, Ojeda F, De Moor B, Moreau Y. Candidate gene prioritization by network analysis of differential expression using machine learning approaches. BMC bioinformatics. 2010 Dec;11(1):1–6. https://doi.org/10.1186/1471-2105-11-460

[5] Glaab E, Bacardit J, Garibaldi JM, Krasnogor N. Using rule-based machine learning for candidate disease gene prioritization and sample classification of cancer gene expression data. PloS one. 2012 Jul 11;7(7):e39932. https://doi.org/10.1371/journal.pone.0039932

[6] Baldi P. Autoencoders, unsupervised learning, and deep architectures. InProceedings of ICML workshop on unsupervised and transfer learning 2012 Jun 27 (pp. 37–49). JMLR Workshop and Conference Proceedings.

[7] Adie EA, Adams RR, Evans KL, Porteous DJ, Pickard BS. SUSPECTS: enabling fast and effective prioritization of positional candidates. Bioinformatics. 2006 Mar 15;22(6):773–4. https://doi.org/10.1093/bioinformatics/btk031

[8] Hutz JE, Kraja AT, McLeod HL, Province MA. CANDID: a flexible method for prioritizing candidate genes for complex human traits. Genetic Epidemiology: The Official Publication of the International Genetic Epidemiology Society. 2008 Dec;32(8):779–90. https://doi.org/10.1002/gepi.20346

[9] Malhotra D, Sebat J. CNVs: harbingers of a rare variant revolution in psychiatric genetics. Cell. 2012 Mar 16;148(6):1223–41. https://doi.org/10.1016/j.cell.2012.02.039

[10] Cheng MC, Chien WH, Huang YS, Fang TH, Chen CH. Translational Study of Copy Number Variations in Schizophrenia. International Journal of Molecular Sciences. 2021 Dec 31;23(1):457. https://doi.org/10.3390/ijms23010457

[11] Coe, Bradley P., et al. “Refining analyses of copy number variation identifies specific genes associated with developmental delay.” Nature genetics 46.10 (2014): 1063–1071. https://doi.org/10.1038/ng.3092

[12] Bromberg Y. Chapter 15: disease gene prioritization. PLoS computational biology. 2013 Apr 25;9(4):e1002902. https://doi.org/10.1371/journal.pcbi.1002902

[13] Tranchevent LC, Ardeshirdavani A, ElShal S, Alcaide D, Aerts J, Auboeuf D, Moreau Y. Candidate gene prioritization with Endeavour. Nucleic acids research. 2016 Jul 8;44(W1):W117–21. https://doi.org/10.1093/nar/gkw365

[14] Stäubert C, Tarnow P, Brumm H, Pitra C, Gudermann T, Gruters A, Schöneberg T, Biebermann H, Römpler H. Evolutionary aspects in evaluating mutations in the melanocortin 4 receptor. Endocrinology. 2007 Oct 1;148(10):4642–8. https://doi.org/10.1210/en.2007-0138

[15] Jiang BB, Wang JG, Wang Y, Xiao J. Gene prioritization for type 2 diabetes in tissue-specific protein interaction networks. Systems Biology. 2009 Sep;10801131:319–28.

[16] Mefford HC, Muhle H, Ostertag P, von Spiczak S, Buysse K, Baker C, Franke A, Malafosse A, Genton P, Thomas P, Gurnett CA. Genome-wide copy number variation in epilepsy: novel susceptibility loci in idiopathic generalized and focal epilepsies. PLoS genetics. 2010 May 20;6(5):e1000962. https://doi.org/10.1371/journal.pgen.1000962

[17] Altman RB, Bergman CM, Blake J, Blaschke C, Cohen A, Gannon F, Grivell L, Hahn U, Hersh W, Hirschman L, Jensen LJ. Text mining for biology-the way forward: opinions from leading scientists. Genome biology. 2008 Sep;9(2):1–5. https://doi.org/10.1186/gb-2008-9-s2-s7

[18] Zolotareva O, Kleine M. A survey of gene prioritization tools for Mendelian and complex human diseases. Journal of integrative bioinformatics. 2019 Dec 1;16(4). https://doi.org/10.1515/jib-2018-0069

[19] Moreau Y, Tranchevent LC. Computational tools for prioritizing candidate genes: boosting disease gene discovery. Nature Reviews Genetics. 2012 Aug;13(8):523–36. https://doi.org/10.1038/nrg3253

[20] Börnigen D, Tranchevent LC, Bonachela-Capdevila F, Devriendt K, De Moor B, De Causmaecker P, Moreau Y. An unbiased evaluation of gene prioritization tools. Bioinformatics. 2012 Dec 1;28(23):3081–8. https://doi.org/10.1093/bioinformatics/bts581

[21] Seyyedrazzagi E, Navimipour NJ. Disease genes prioritizing mechanisms: a comprehensive and systematic literature review. Network Modeling Analysis in Health Informatics and Bioinformatics. 2017 Dec;6(1):1–5. https://doi.org/10.1007/s13721-017-0154-9

[22] Goodman SN. Toward evidence-based medical statistics. 1: The P value fallacy. Annals of internal medicine. 1999 Jun 15;130(12):995–1004. https://doi.org/10.7326/0003-4819-130-12-199906150-00008

[23] Boudellioua I, Kulmanov M, Schofield PN, Gkoutos GV, Hoehndorf R. DeepPVP: phenotype-based prioritization of causative variants using deep learning. BMC bioinformatics. 2019 Dec;20(1):1–8. https://doi.org/10.1101/311621

[24] Zakeri P, Simm J, Arany A, ElShal S, Moreau Y. Gene prioritization using Bayesian matrix factorization with genomic and phenotypic side information. Bioinformatics. 2018 Jul 1;34(13):i447–56. https://doi.org/10.1093/bioinformatics/bty289

[25] Kingma DP, Welling M. Auto-encoding variational Bayes. arXiv preprint arXiv:1312.6114. 2013 Dec 20.

[26] Kingma D, Welling M. Efficient gradient-based inference through transformations between Bayes nets and neural nets. In International Conference on Machine Learning 2014 Jun 18 (pp. 1782–1790). PMLR.

[27] Firth HV, Richards SM, Bevan AP, Clayton S, Corpas M, Rajan D, Van Vooren S, Moreau Y, Pettett RM, Carter NP. DECIPHER: database of chromosomal imbalance and phenotype in humans using ensemble resources. The American Journal of Human Genetics. 2009 Apr 10;84(4):524–33. https://doi.org/10.1016/j.ajhg.2009.03.010

[28] Alinejad-Rokny H, Heng JI, Forrest AR. Brain-enriched coding and long non-coding RNA genes are overrepresented in recurrent neurodevelopmental disorder CNVs. Cell Reports. 2020 Oct 27;33(4):108307. https://doi.org/10.1016/j.celrep.2020.108307

[29] Marshall CR, Howrigan DP, Merico D, Thiruvahindrapuram B, Wu W, Greer DS, Antaki D, Shetty A, Holmans PA, Pinto D, Gujral M. Contribution of copy number variants to schizophrenia from a genome-wide study of 41,321 subjects. Nature genetics. 2017 Jan;49(1):27–35. https://doi.org/10.1038/ng.3725

[30] The Remap Tool. https://www.ncbi.nlm.nih.gov/genome/tools/remap

[31] Pinto D, Delaby E, Merico D, Barbosa M, Merikangas A, Klei L, Thiruvahindrapuram B, Xu X, Ziman R, Wang Z, Vorstman JA. Convergence of genes and cellular pathways dysregulated in autism spectrum disorders. The American Journal of Human Genetics. 2014 May 1;94(5):677–94. https://doi.org/10.1016/j.ajhg.2014.03.018

[32] Hon CC, Ramilowski JA, Harshbarger J, Bertin N, Rackham OJ, Gough J, Denisenko E, Schmeier S, Poulsen TM, Severin J, Lizio M. An atlas of human long non-coding RNAs with accurate 5′ ends. Nature. 2017 Mar;543(7644):199–204. https://doi.org/10.1038/nature21374

[33] Cooper GM, Coe BP, Girirajan S, Rosenfeld JA, Vu TH, Baker C, Williams C, Stalker H, Hamid R, Hannig V, Abdel-Hamid H. A copy number variation morbidity map of developmental delay. Nature genetics. 2011 Sep;43(9):838–46. https://doi.org/10.1038/ng.909

[34] Purcell S, Neale B, Todd-Brown K, Thomas L, Ferreira MA, Bender D, Maller J, Sklar P, De Bakker PI, Daly MJ, Sham PC. PLINK: a tool set for whole-genome association and population-based linkage analyses. The American journal of human genetics. 2007 Sep 1;81(3):559–75. https://doi.org/10.1086/519795

[35] Liao Y, Wang J, Jaehnig EJ, Shi Z, Zhang B. WebGestalt 2019: gene set analysis toolkit with revamped UIs and APIs. Nucleic acids research. 2019 Jul 2;47(W1): W199–205. https://doi.org/10.1093/nar/gkz401

[36] Ashburner M, Ball CA, Blake JA, Botstein D, Butler H, Cherry JM, Davis AP, Dolinski K, Dwight SS, Eppig JT, Harris MA. Gene ontology: tool for the unification of biology. Nature genetics. 2000 May;25(1):25–9. https://doi.org/10.1038/75556

[37] Piñero J, Bravo À, Queralt-Rosinach N, Gutiérrez-Sacristán A, Deu-Pons J, Centeno E, García-García J, Sanz F, Furlong LI. DisGeNET: a comprehensive platform integrating information on human disease-associated genes and variants. Nucleic acids research. 2016 Oct 19:gkw943. https://doi.org/10.1093/nar/gkw943

[38] Fallah MS, Eubanks JH. Seizures in mouse models of rare neurodevelopmental disorders. Neuroscience. 2020 Oct 1;445:50–68. https://doi.org/10.1016/j.neuroscience.2020.01.041

[39] Cardoso AR, Lopes-Marques M, Silva RM, Serrano C, Amorim A, Prata MJ, Azevedo L. Essential genetic findings in neurodevelopmental disorders. Human genomics. 2019 Dec;13(1):1–7. https://doi.org/10.1186/s40246-019-0216-4

[40] Kingma DP, Mohamed S, Jimenez Rezende D, Welling M. Semi-supervised learning with deep generative models. Advances in neural information processing systems. 2014;27.

[41] DGT RP, FANTOM Consortium. A promoter-level mammalian expression atlas. Nature. 2014 Mar 27;507(7493):462–70. https://doi.org/10.1038/nature13182

[42] Bult CJ, Eppig JT, Kadin JA, Richardson JE, Blake JA, Mouse Genome Database Group. The Mouse Genome Database (MGD): mouse biology and model systems. Nucleic acids research. 2008 Jan;36(suppl_1): D724–8. https://doi.org/10.1093/nar/gkm961

[43] Mouse 0003631. http://www.informatics.jax.org/mp/annotations/MP:0003631

[44] Mouse 0003632. http://www.informatics.jax.org/mp/annotations/MP:0003632

[45] Mouse 0003633. http://www.informatics.jax.org/mp/annotations/MP:0003633

[46] Mouse Homologene. http://www.informatics.jax.org/downloads/reports/HGNChomologene.rpt

[47] Kent WJ, Sugnet CW, Furey TS, Roskin KM, Pringle TH, Zahler AM, Haussler D. The human genome browser at UCSC. Genome research. 2002 Jun 1;12(6):996–1006. https://doi.org/10.1101/gr.229102

[48] Chicco D, Sadowski P, Baldi P. Deep autoencoder neural networks for gene ontology annotation predictions. InProceedings of the 5th ACM conference on bioinformatics, computational biology, and health informatics 2014 Sep 20 (pp. 533–540). https://doi.org/10.1145/2649387.2649442

[49] Chen L, Cai C, Chen V, Lu X. Learning a hierarchical representation of the yeast transcriptomic machinery using an autoencoder model. InBMC bioinformatics 2016 Dec (Vol. 17, No. 1, pp. 97–107). BioMed Central. https://doi.org/10.1186/s12859-015-0852-1

[50] Svensson V, Gayoso A, Yosef N, Pachter L. Interpretable factor models of single-cell RNA-seq via variational autoencoders. Bioinformatics. 2020 Jun 1;36(11):3418–21. https://doi.org/10.1093/bioinformatics/btaa169

[51] Doersch C. Tutorial on variational autoencoders. arXiv preprint arXiv:1606.05908. 2016 Jun 19.

[52] Genevay A, Peyré G, Cuturi M. GAN and VAE from an optimal transport point of view. arXiv preprint arXiv:1706.01807. 2017 Jun 6.

[53] May T, Adesina I, McGillivray J, Rinehart NJ. Sex differences in neurodevelopmental disorders. Current opinion in neurology. 2019 Aug 1;32(4):622–6. https://doi.org/10.1097/WCO.0000000000000714

[54] Rinehart NJ, Cornish KM, Tonge BJ. Gender differences in neurodevelopmental disorders: Autism and fragile x syndrome. Biological basis of sex differences in psychopharmacology. 2010:209–29. https://doi.org/10.1007/7854_2010_96

[55] Brentani H. Gender, Genetic, And Environmental Factors In The Neurodevelopmental Disorders. European Neuropsychopharmacology. 2019 Jan 1;29:S745–6. https://doi.org/10.1016/j.euroneuro.2017.06.083

[56] Al-Beltagi M. Autism medical comorbidities. World journal of clinical pediatrics. 2021 May 9;10(3):15. https://doi.org/10.5409/wjcp.v10.i3.15

[57] Buckley PF, Miller BJ, Lehrer DS, Castle DJ. Psychiatric comorbidities and schizophrenia. Schizophrenia bulletin. 2009 Mar 1;35(2):383–402. https://doi.org/10.1093/schbul/sbn135

[58] Schilbach L. Autism and other disorders of social interaction: where we are and where to go from here. European Archives of Psychiatry and Clinical Neuroscience. 2022 Feb 9:1–3. https://doi.org/10.1007/s00406-022-01391-y

[59] Hisaoka T, Komori T, Kitamura T, Morikawa Y. Abnormal behaviors relevant to neurodevelopmental disorders in Kirrel3-knockout mice. Scientific reports. 2018 Jan 23;8(1):1–2. https://doi.org/10.1038/s41598-018-19844-7

[60] Martínez-Cerdeño V. Dendrite and spine modifications in autism and related neurodevelopmental disorders in patients and animal models. Developmental neurobiology. 2017 Apr;77(4):393–404. https://doi.org/10.1002/dneu.22417

[61] Zieger HL, Choquet D. Nanoscale synapse organization and dysfunction in neurodevelopmental disorders. Neurobiology of Disease. 2021 Oct 1;158:105453. https://doi.org/10.1016/j.nbd.2021.105453

[62] Xue, Chen, et al. “Progress and assessment of lncRNA DGCR5 in malignant phenotype and immune infiltration of human cancers.” American Journal of Cancer Research 11.1 (2021): 1.

[63] Suzuki G, Harper KM, Hiramoto T, Sawamura T, Lee M, Kang G, Tanigaki K, Buell M, Geyer MA, Trimble WS, Agatsuma S. Sept5 deficiency exerts pleiotropic influence on affective behaviors and cognitive functions in mice. Human molecular genetics. 2009 May 1;18(9):1652–60. https://doi.org/10.1093/hmg/ddp086

[64] Molinard-Chenu A, Dayer A. The candidate schizophrenia risk gene DGCR2 regulates early steps of corticogenesis. Biological Psychiatry. 2018 Apr 15;83(8):692–706. https://doi.org/10.1016/j.biopsych.2017.11.015

[65] Hyde TM, Weinberger DR. Seizures and schizophrenia. Schizophrenia bulletin. 1997 Jan 1;23(4):611–22. https://doi.org/10.1093/schbul/23.4.611

[66] Kunugi H, Takei N, Murray RM, Saito K, Nanko S. Small head circumference at birth in schizophrenia. Schizophrenia research. 1996 May 1;20(1-2):165–70. https://doi.org/10.1016/0920-9964(96)00007-2

[67] Klein S, Sharifi-Hannauer P, Martinez-Agosto JA. Macrocephaly as a clinical indicator of genetic subtypes in autism. Autism Research. 2013 Feb;6(1):51–6. https://doi.org/10.1002/aur.1266

[68] Tripi G, Roux S, Matranga D, Maniscalco L, Glorioso P, Bonnet-Brilhault F, Roccella M. Cranio-facial characteristics in children with autism spectrum disorders (ASD). Journal of Clinical Medicine. 2019 May 9;8(5):641. https://doi.org/10.3390/jcm8050641

[69] Hosseini MP, Beary M, Hadsell A, Messersmith R, Soltanian-Zadeh H. Deep Learning for Autism Diagnosis and Facial Analysis in Children. Frontiers in Computational Neuroscience. 2021;15. https://doi.org/10.3389/fncom.2021.789998

[70] Chourasia N, Ossó-Rivera H, Ghosh A, Von Allmen G, Koenig MK. Expanding the phenotypic spectrum of CACNA1H mutations. Pediatric Neurology. 2019 Apr 1;93:50–5. https://doi.org/10.1016/j.pediatrneurol.2018.11.017

[71] Torti E, Keren B, Palmer EE, Zhu Z, Afenjar A, Anderson I, Andrews MV, Atkinson C, Au M, Berry SA, Bowling KM. Variants in TCF20 in neurodevelopmental disability: description of 27 new patients and review of literature. Genetics in Medicine. 2019 Sep;21(9):2036–42. https://doi.org/10.1038/s41436-019-0454-9

[72] Xie B, Fan X, Lei Y, Chen R, Wang J, Fu C, Yi S, Luo J, Zhang S, Yang Q, Chen S. A novel de novo microdeletion at 17q11. 2 adjacent to NF1 gene associated with developmental delay, short stature, microcephaly and dysmorphic features. Molecular cytogenetics. 2016 Dec;9(1):1–5. https://doi.org/10.1186/s13039-016-0251-y

